# Parental Genome Assemblies of Suyunuo1 Reveal Structural Variation Underlying Edible Waxy Maize Evolution, Superior Hybrid Performance and Yield - Flavor Balance

**DOI:** 10.64898/2026.09.24.754239

**Authors:** Ling Zhou, Jun Hong, Wenming Zhao, Tifu Zhang, Lihua Ning, Long Ruan, Huixue Dai, Han Zhao

## Abstract

Edible waxy maize is a unique domesticated cereal valued for its superior sensory and nutritional properties, yet high-quality gap-free genomes remain lacking, hindering the exploration of structural variations (SVs) and domestication-related divergence. Here, we assembled two chromosome-level, gap-free genomes of the elite waxy maize inbred lines Tongxi 5 and Hengbai 522, the parents of the widely cultivated hybrid Suyunuo 1. Comparative genomic analysis revealed substantial genome size variation, prominent megabase-scale SVs, and extensive knob-repeat expansion in Tongxi 5. We identified widespread gene presence–absence variation and hyperdivergent regions enriched in transposable elements, which largely underpin genomic differentiation between waxy and field maize. Population genomic analyses further demonstrated asymmetric introgression from field maize into the two parents, whereas the conserved waxy haplotype supports a shared ancestral origin. Integrative GWAS based on SNPs, InDels and SVs revealed that SVs substantially contribute to phenotypic diversity and mediate the co-regulation of yield traits and flavor-related metabolites. Our results uncover key genomic events underlying post-domestication divergence and highlight the essential roles of SVs in heterosis and flavor–yield balance, providing valuable genomic resources for waxy maize improvement.

---

Edible waxy maize is a distinct cultivated type that emerged through mutation at the *waxy* locus and subsequent post-domestication evolution after non-glutinous field maize was introduced from the New World ∼500 years ago (Li et al., 2025a). Valued for its distinctive sensory attributes and nutritional profile, edible waxy maize has long been cultivated as both a vegetable and staple food across much of Asia, where glutinous foods are deeply embedded in traditional diets and culinary heritage. Despite advances from short-read resequencing (Li et al., 2025a; Luo et al., 2024), the lack of a high-quality waxy maize reference genome has limited resolution of large structural variants and complex, repeat-rich regions relevant to adaptation and improvement. Suyunuo 1, the first nationally approved edible waxy maize variety in China, combines pronounced heterosis with exceptional flavor quality and has contributed to the development of several elite second-cycle inbred lines, including JS118, Zhengbainuo 04, CTW3446 and SU-2. Gapless genome assemblies of the two parental lines, Tongxi 5 and Hengbai 522 (Figure 1a, Figure S1), therefore provide a foundation for investigating edible waxy maize germplasm innovation, expand the genomic representation of maize diversity and enable comparative genomic analyses.

**Figure 1.**
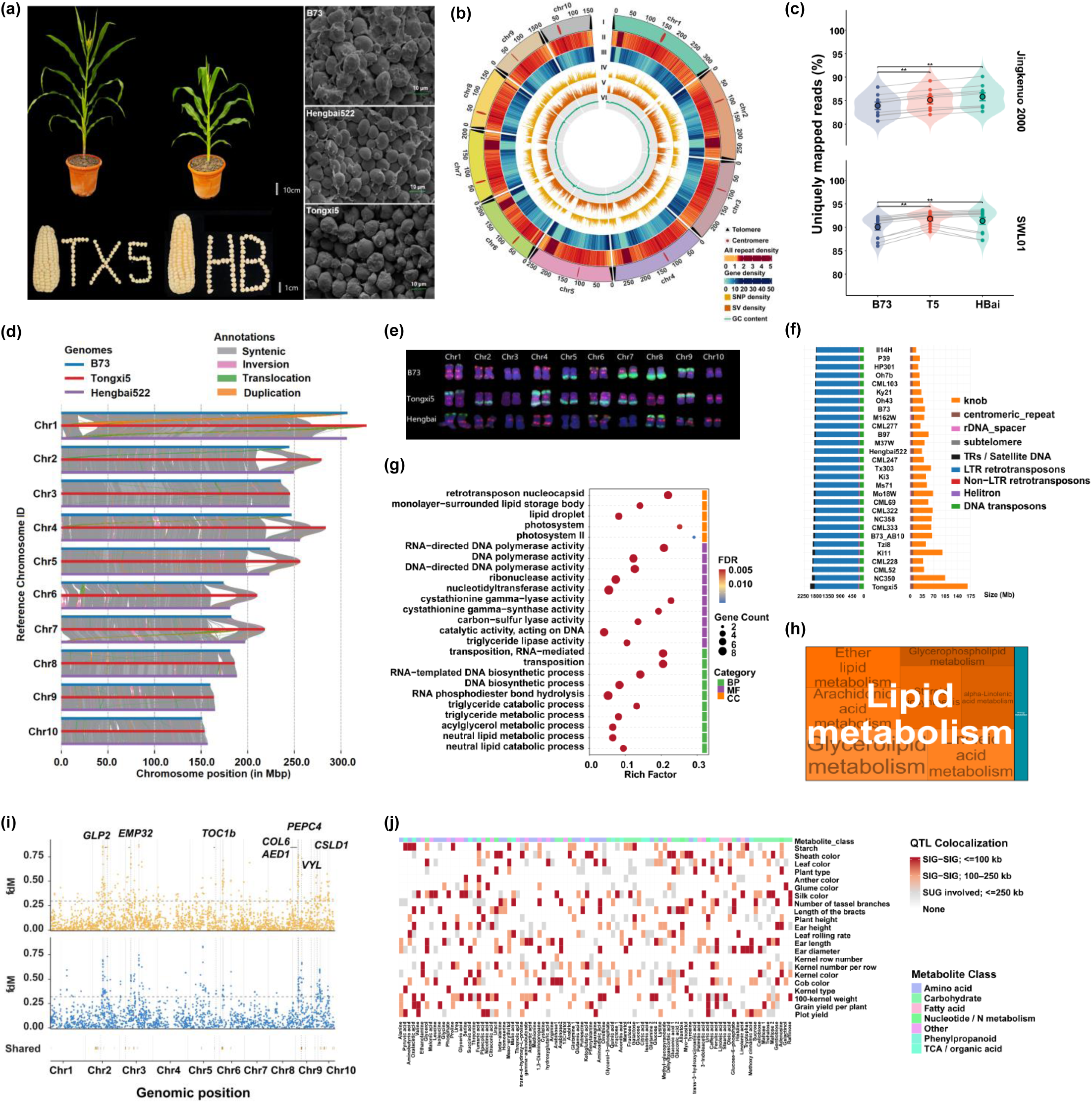
Genome features, comparative genomics and introgression in edible waxy maize. (a) Plant architecture and starch granule morphology of the two parental lines. (b) Circular view of the Tongxi 5 genome showing the genome-wide distribution of major genomic features across the ten chromosomes. From the outermost to the innermost tracks of the Circos plot: (I) telomere and centromere; (II) repeat density (number of repeats per 500 kb); (III) gene density (number of genes per 500 kb); (IV) SNP density (number of SNPs per 500 kb); (V) structural variation density (number of SVs per 500 kb); and (VI) GC content calculated in 500-kb windows. Telomeres are indicated by black triangles and centromeres in red. (c) Uniquely mapped RNA-seq read percentages for Jingkenuo 2000 (n = 9) and SWL01 (n = 12) against the B73, T5 and HBai genomes; points indicate samples, grey lines paired measurements and large circles means ± s.e.m. Significance was assessed by two-sided paired Wilcoxon signed-rank tests. (d) Synteny and SVs among B73, Tongxi 5 and Hengbai 522. (e) FISH-based karyotypes of B73, Tongxi 5 and Hengbai 522. (f) Comparison of transposable element and repetitive sequence content across maize genomes. Bar lengths represent the total genomic sequence length (Mb) occupied by each repeat class in each genome. (g) GO enrichment of shared-specific reciprocal best-hit (RBH) genes. Point size represents gene count, point color denotes FDR and the x axis indicates Rich Factor. Colored bars indicate biological process (green), molecular function (purple) and cellular component (orange). (h) KEGG enrichment of the shared-specific genes. Significantly enriched pathways (FDR < 0.05) are displayed as a treemap, with rectangle area proportional to gene count × −log10 (FDR). Pathways are grouped by functional category, with related pathways shown in different shades of the same color. (i) Genome-wide introgression signals in Hengbai 522. Positive f_dM values from ABBA-BABAwindows (upper) and Dsuite Dinvestigate (middle) indicate excess allele sharing between Hengbai 522 and field maize. Dashed horizontal lines denote the 90th percentile of positive values used to define high-signal candidate windows, and grey shading marks genomic intervals supported by both methods. Dotted vertical lines indicate key genes located within shared introgression regions. The lower track summarizes shared candidate introgression intervals across chromosomes. (j) Agronomic trait–metabolite GWAS colocalization heatmap. Rows denote agronomic traits and columns metabolites; Local scores indicate colocalization strength within 250 kb (0, none; 1, ≥1 suggestive association; 2, significant–significant pairs at 100–250 kb; 3, significant–significant pairs within 100 kb). Color intensity reflects increasing colocalization strength, and the top annotation indicates metabolite class.

Here, we generated four types of high coverage sequencing datasets for the assembly of two waxy inbred lines (Tongxi 5 and Hengbai 522), comprising PacBio HiFi (151 Gb / 86 Gb), Nanopore ultra long (175 Gb / 203 Gb), short-read paired-end (88 Gb/ 117 Gb), and Hi C data (103 Gb / 121 Gb). We assembled two gap-free chromosome-level genomes for Tongxi 5 (2324.54 Mb) and Hengbai 522 (2168.39 Mb), respectively (Figure 1b, Figure S2), with contig N50 values of 186.09 Mb/188.39 Mb, scaffold N50 values of 245.35 Mb/223.07 Mb, BUSCO completeness of 98.2%/97.9%, LTR Assembly Index scores of 24.95/27.39, and Merqury QV socres of 43.92/43.09 (Figure S3 and S4). Only one chromosome end in each assembly lacked an identifiable telomere, on chromosomes 4 and 2 in Tongxi 5 and Hengbai 522, respectively (Figure 1b, Figure S2). Repetitive sequences comprised 88.29% and 87.40% of the respective genomes, and gene annotation identified 41,468 and 42,324 protein-coding genes, with BUSCO completeness of 98.8% and 99.1%. Together, these metrics demonstrate the high contiguity, consensus accuracy and completeness of both assemblies. Given that reference-genome choice can influence transcriptome profiling (Igolkina et al., 2025), we reanalyzed 21 samples from two published waxy maize RNA-seq datasets against the Tongxi 5, Hengbai 522 and B73 genomes. Both waxy maize assemblies yielded higher proportions of uniquely mapped reads than B73 (Figure 1c), indicating improved representation of waxy maize transcriptomes and supporting their utility as reference resources.

Notably, the Tongxi 5 genome assembly (2,324.54 Mb) was ∼7.2% larger than that of Hengbai 522 (2,168.39 Mb). Comparative collinearity analysis revealed extensive genomic divergence between the two lines, with syntenic regions covering only 1.36 Gb (58.6% of the Tongxi 5 genome). Six prominent megabase-scale structural variations were detected in Tongxi 5 on chromosomes 1, 2, 4, 5, 6 and 7, ranging from 20.8 to 35.1 Mb and collectively spanning 172.9 Mb (Figure 1d). EDTA-based annotation showed that Tongxi 5-specific expanded regions were strongly enriched for knob-associated repeats relative to the corresponding regions in Hengbai 522 and B73. Their presence and chromosomal distribution were further supported by fluorescence in situ hybridization (FISH) with knob-specific probes and PacBio read alignment (Figure 1e, Figure S5). Genome-wide repeat profiling across Tongxi 5 and 26 NAM founder lines (Hufford et al., 2021) revealed a greater abundance of knob-associated repeats in Tongxi 5 than in any other line examined (Figure 1f), suggesting that this expansion reflects its unique post-domestication evolutionary trajectory and extending the contribution of knob repeats to genome-size variation from field maize to waxy maize germplasm (Tan et al., 2026).

Comparative pan-genome analysis of the two parental genomes and 26 NAM founders revealed extensive gene presence–absence variation, providing additional insight into field–waxy genomic differentiation beyond that captured by short-read resequencing (Li et al., 2025a; Luo et al., 2024). Tongxi 5 and Hengbai 522 contained 4,045 and 4,414 genes, respectively, without detectable homologs in field maize, including 1,900 and 2,036 parent-specific genes and 1,259 reciprocal best-hit gene pairs shared by both parents. These shared waxy lineage-specific genes were enriched for lipid-related pathways, including glycerolipid, glycerophospholipid, linoleic acid and α-linolenic acid metabolism (Tables S1 and S2, Figures 1g and 1h), revealing unresolved field–waxy divergence in kernel lipid homeostasis and quality-related metabolism. Hyperdivergent regions (HDRs), characterized by low sequence homology and high variant density, are often underrepresented in conventional population genomic analyses yet contribute substantially to maize population differentiation (Feng et al.). We compared HDR landscapes between the two parental lines and B73 to assess their role in waxy–field genomic differentiation. Tongxi 5 and Hengbai 522 contained 21,772 and 21,016 HDRs spanning 376.0 and 403.6 Mb, respectively, with 149.8 Mb shared and 226.2 Mb and 253.8 Mb specific to each parent (Figures S6 and S7). Notably, 69.4–71.9% of HDR sequences overlapped transposable elements, predominantly Gypsy/Copia LTR retrotransposons and helitrons (Figure S8), linking parental genomic divergence to repeat-rich regions. Shared HDRs contained an ERF9-like GC-rich motif and a DOF3.4-like T-rich motif. Tongxi5-specific HDRs showed a similar ERF9-like GC-rich motif together with a CDF5-like T-rich motif, whereas Hengbai522-specific HDRs were characterized by an AS2-like GC-rich motif and a CDF5-like T-rich motif (Figure S9). These patterns indicate divergence in HDR boundary-associated cis-regulatory features, potentially reflecting distinct regulatory architectures between the parental genomes.

Although field and waxy maize are genetically differentiated, gene flow between them remains poorly understood. Resolving introgression in waxy maize parental lines could clarify their ancestry and its contribution to heterosis during crop improvement (Lin et al., 2020). Resequencing of 185 maize inbred lines (105 field and 80 waxy) against B73 RefGen_v4 identified 6.88 million SNPs and 467,232 InDels (Table S3). Principal component analysis separated field and waxy maize into distinct genetic groups, with moderate genome-wide population differentiation (genome-wide mean FST*_Field-Waxy_* = 0.10583; Table S4). Within this structure, Tongxi 5 was more strongly differentiated from field maize than Hengbai 522 (Figure S10), consistent with its local waxy landrace origin and suggesting greater field-maize introgression in Hengbai 522. Patterson’s D statistic confirmed significantly greater allele sharing between Hengbai 522 and field maize than between Tongxi 5 and field maize (D = 0.0741, Z = 3.39, P = 7.05 × 10, Figure S11). Concordant ABBA-BABAwindows and Dsuite Dinvestigate analyses further identified 38 candidate introgressed regions spanning ∼57.4 Mb across seven chromosomes (Figure 1i, Table S5), encompassing genes associated with photosynthesis, photoperiod sensitivity, plant architecture and seed development, including *VYL*, *PEPC4*, *TOC1b*, *COL6*, *CSLD1*, *GLP2*, *EMP32* and *AED1*. Together, these results indicate that asymmetric field-maize introgression contributed to post-domestication divergence between the two parental lines. Against this genome-wide divergence, the 300-kb region surrounding the *waxy* locus showed a 98.77% reduction in variant density relative to the genomic background (Welch’s t-test, P < 0.001; Figures S12 and S13). The identical ∼174-kb segment flanking *waxy* in Hengbai 522 and Tongxi 5 is unlikely to have arisen independently, providing strong evidence for a common ancestral origin. Taken together, these findings support two possible scenarios: the parental lines either share a common ancestry followed by independent divergence and differential introgression, or diverged earlier and subsequently acquired the same *waxy* haplotype through breeding-mediated introgression.

Balancing grain yield and flavor quality remains a major challenge in edible waxy maize breeding. To dissect their genetic links, we performed parallel SNP/InDel-and SV-based GWAS for 22 agronomic traits and 91 flavor-related metabolites in 177 maize inbred lines. We detected 3,734 QTN associations, comprising 2,759 SNP/InDel-and 975 SV-based associations and representing 3,625 non-redundant loci (Figure S14, Table S6). Of these, 2,886 were significant, including 2,267 SNP/InDel and 619 SV loci, whereas 739 were suggestive (439 SNP/InDel and 300 SV loci), demonstrating the complementary contribution of SVs to agronomic and metabolic variation. Among significant loci with comparable parental genotypes, 28.59% of SNP/InDel loci (578/2,022) but 82.00% of SV loci (483/589) differed between Tongxi 5 and Hengbai 522 (Table S7), indicating substantially greater parental differentiation at SV-associated loci and considerable potential for allelic complementation in Suyunuo 1. We further identified 595 agronomic–metabolic QTN pairs within 250 kb, of which 394 were significant for both traits, including 188 high-confidence pairs separated by ≤100 kb (Figure 1j, Figure S15). Clustering these dual-significant signals defined 245 candidate QTL regions, 97 of which were supported by both SNP/InDel and SV associations (Table S8). Notable examples included an SV at 123.48 Mb on chromosome 7 associated with both tassel branch number and sucrose, and a chromosome 1 hotspot at 103.67–103.74 Mb linking grain yield per plant and hundred-grain weight with oxaloacetic acid. Notably, several yield–metabolite regions were supported exclusively by SV associations. An SV-only interval on chromosome 5 linked kernel number per row with cysteine, a precursor of sulfur-containing aroma compounds, whereas a chromosome 6 interval connected the same trait with ferulic acid, a phenolic compound associated with bitterness and astringency. Most strikingly, the same SV on chromosome 10 was associated with both ear height and adenosine, a metabolite implicated in sweet-taste signaling. Together, these colocalized associations reveal a partially shared genetic basis of yield productivity and flavor metabolism and highlight SV-rich genomic regions as potential targets for jointly improving yield and eating quality.

Although numerous field-maize genomes have been sequenced, genomic resources for specialized kernel types such as waxy maize remain limited. Long-term artificial selection and local adaptation have shaped waxy maize into a genomically distinct form relative to field maize, underscoring its value as a reservoir of maize genetic diversity (Fang et al., 2025; Li et al., 2025a; Li et al., 2025b). The near-complete parental assemblies generated here address a critical gap in reference resources for edible waxy maize and provide a foundation for favorable-allele discovery and comparative genomics. Our assembly-based analyses uncovered post-domestication divergence poorly captured by short-read resequencing, including knob-repeat expansion, gene-content variation and HDRs, while population genomics revealed asymmetric field-maize introgression into the two parental lines. We also highlighted the contribution of structural variation to heterosis and to the balance between yield and flavor metabolism in edible waxy maize. Together, these findings provide a genomic framework for understanding edible waxy maize evolution, germplasm innovation and the genetic basis of parental complementation in Suyunuo 1.

## AUTHOR CONTRIBUTIONS

Han Zhao conceived, supervised the project, obtained funding and revised the manuscript. Ling Zhou and Jun Hong contributed equally: Ling Zhou completed genome assembly, annotation, transcriptome and pan-genome analyses; Jun Hong performed population genomics, introgression and GWAS analyses. Wenming Zhao assisted comparative genomics and functional enrichment. Long Ruan and Huixue Dai conducted field phenotyping, sample collection and metabolomic preprocessing. Tifu Zhang and Lihua Ning supported sequencing-data processing and genome quality assessment. All authors read and approved the final manuscript.

## ACKNOWLEDGMENTS

We thank the staff at the Maize Research Center, Jiangsu Academy of Agricultural Sciences for field-related technical support. We appreciate Guangdong Academy of Agricultural Sciences for providing metabolome testing service.

## FUNDING

This work was supported by the Agricultural Science and Technology Independent Innovation Fund Project of Jiangsu Province (CX [24]3090), and the National Natural Science Foundation of China (32272133), (32372101).

## COMPETING INTERESTS

The authors declare no competing interests, and approved the paper.

## DATA AVAILABILITY STATEMENT

Genomic data supporting the findings of this study are available in the NCBI database (https://www.ncbi.nlm.nih.gov/sra) under the BioProject accession PRJNA1523807, PRJNA1523877, PRJNA1524114, PRJNA1524281, PRJNA1524485 and PRJNA1525083. The above raw sequencing data are Hi C, PacBio HiFi, Illumina short read whole genome sequencing data, RNA seq transcriptome sequencing data, Oxford Nanopore whole genome sequencing data of two parental lines, and whole genome sequencing data of 185 waxy maize inbred lines. The de novo assembled genomes and annotation, PRJNA1524947.

## SHORT LEGENDS FOR SUPPORTING INFORMATION

### Supporting Information

**Figure S1.**
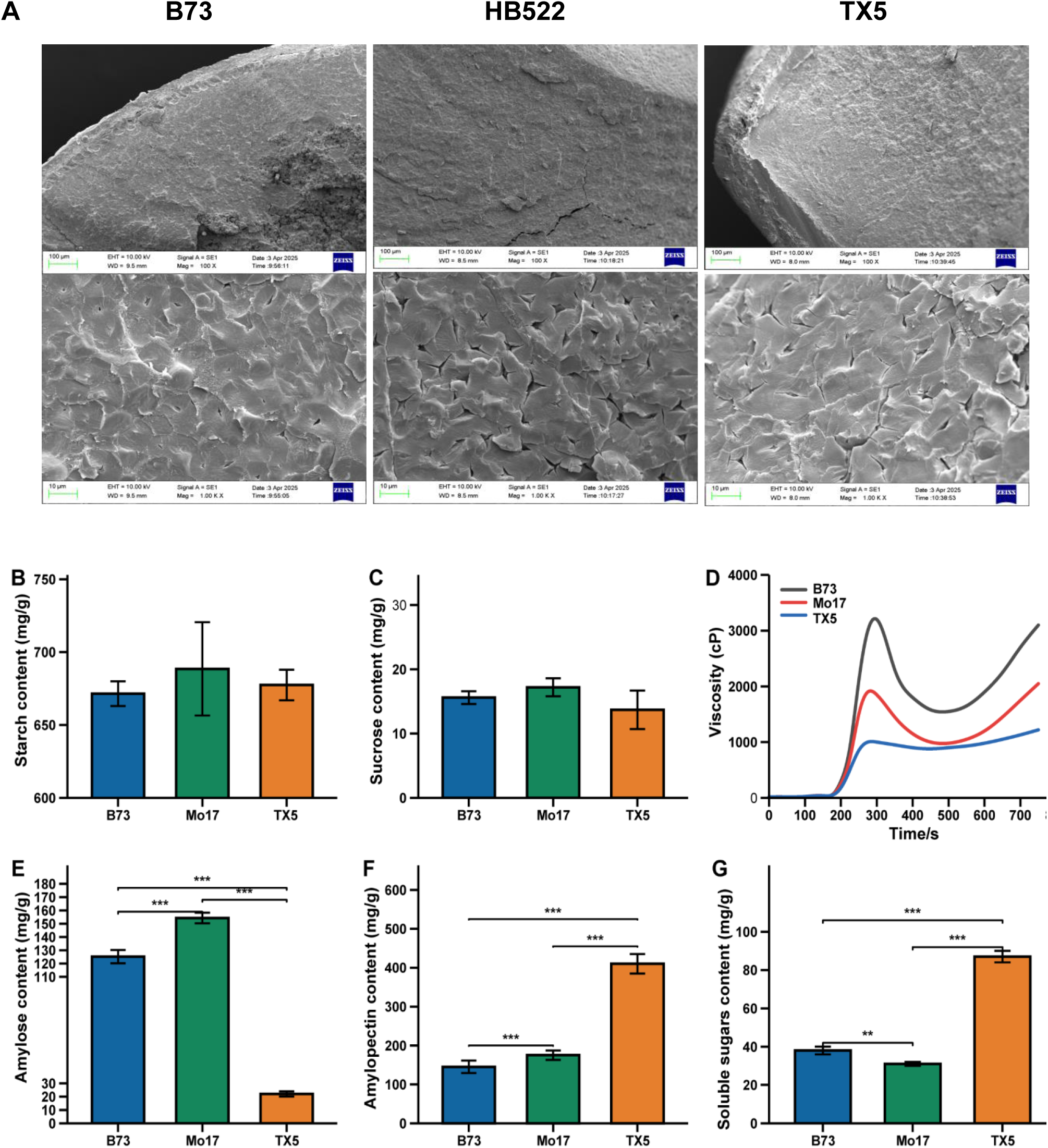
Starch granule morphology, carbohydrate composition and pasting properties of maize kernels. **(A)** Representative scanning electron microscopy (SEM) images of starch granules from B73, Hengbai 522 (HB522) and Tongxi 5 (TX5). **(B)** Starch content, **(C)** sucrose content, **(D)** viscosity profiles, **(E)** amylose content, **(F)** amylopectin content and **(G)** soluble sugar content in B73, Mo17, TX5. Bars represent mean ± SE. Horizontal brackets indicate pairwise statistical comparisons; *P < 0.05, **P < 0.01 and ***P < 0.001, representing significant, highly significant and extremely highly significant differences, respectively.

**Figure S2.**
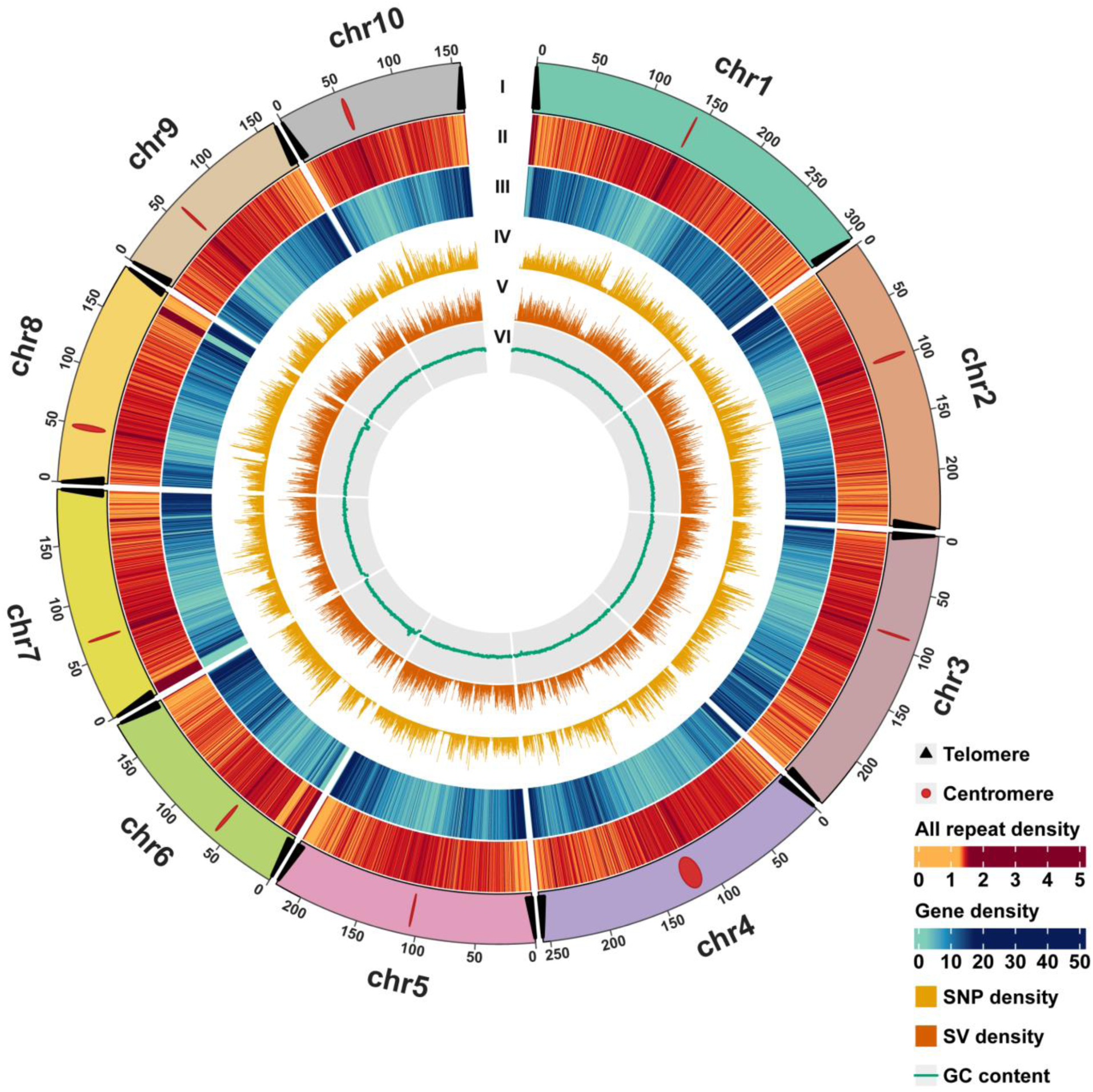
Circular view of the Hengbai522 genome showing the genome-wide distribution of major genomic features across the ten chromosomes. From the outermost to the innermost tracks of the Circos plot: (I) telomere and centromere positions; (II) repeat density (number of repeats per 500 kb); (III) gene density (number of genes per 500 kb); (IV) SNP density (number of SNPs per 500 kb); (V) structural variation (SV) density (number of SVs per 500 kb); and (VI) GC content calculated in 500-kb windows. Chromosome coordinates are shown in megabases (Mb). Telomeres are indicated by black triangles and centromeres in red.

**Figure S3.**
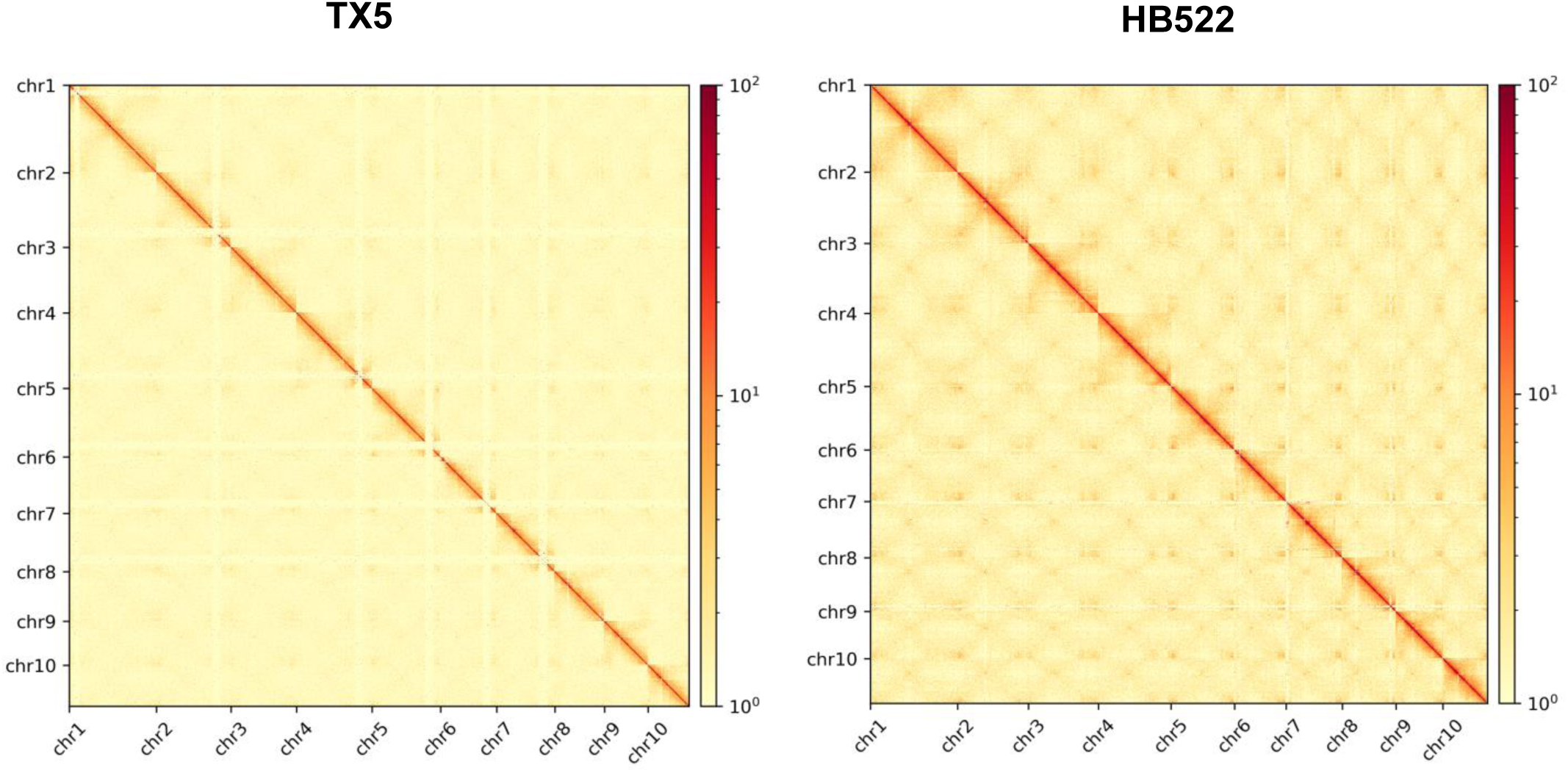
Genome-wide Hi-C interaction maps of Tongxi 5 (TX5) and Hengbai 522 (HB522) across the ten chromosomes. Color intensity represents interaction frequency on a logarithmic scale, with stronger contacts shown in darker colors. Continuous diagonal interaction patterns support the chromosome-scale organization of both assemblies.

**Figure S4.**
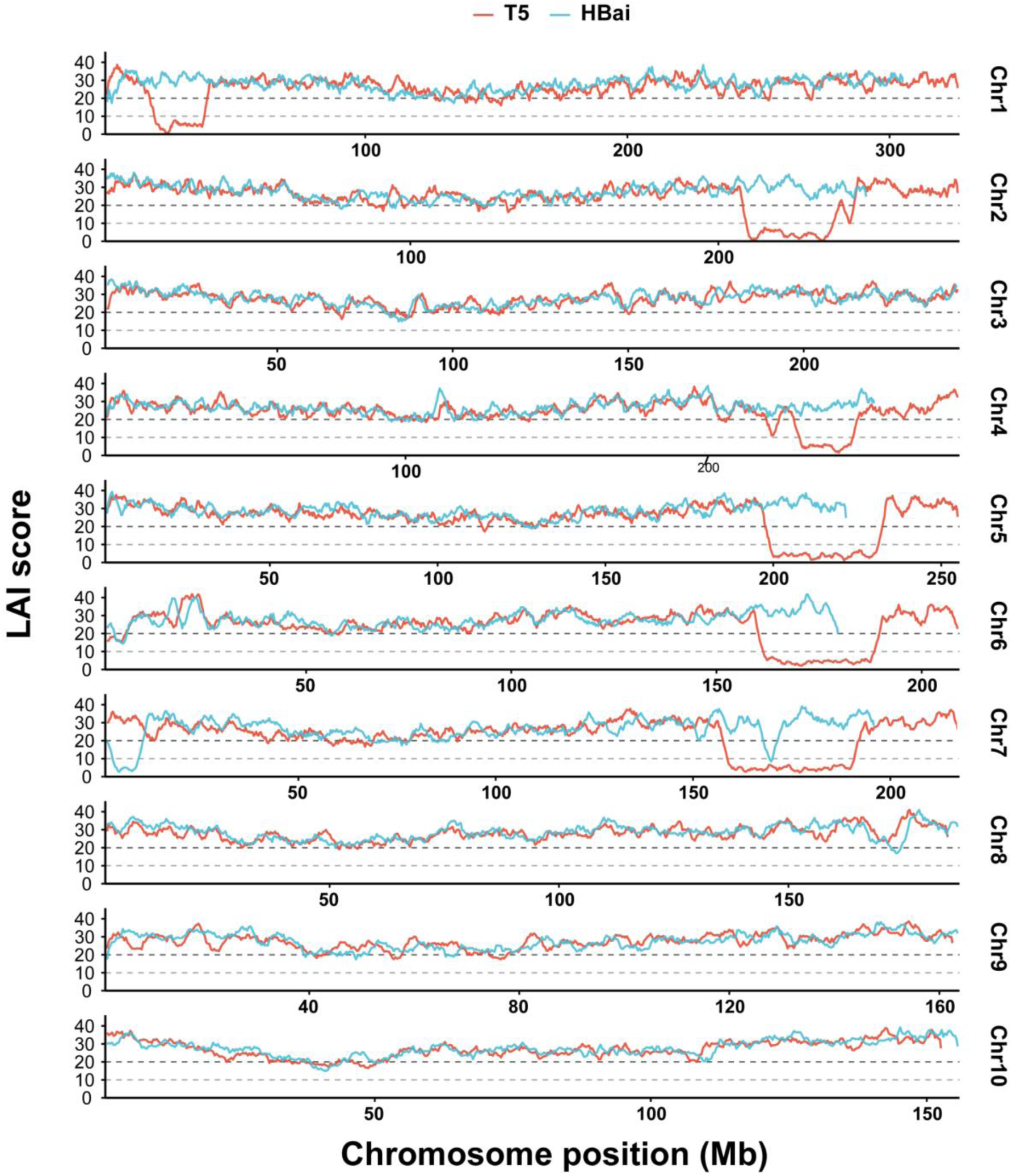
Genome-wide LTR Assembly Index (LAI) profiles of the Tongxi5 (T5) and Hengbai522 (HBai) genomes. Local LAI scores were calculated across the ten chromosomes using 3-Mb sliding windows with a 300-kb step. T5 and HBai are shown in red and blue, respectively. Chromosomal positions are shown in megabases (Mb). Dashed horizontal lines indicate LAI scores of 10 and 20, providing reference thresholds for evaluating local assembly continuity in repeat-rich genomic regions.

**Figure S5.**
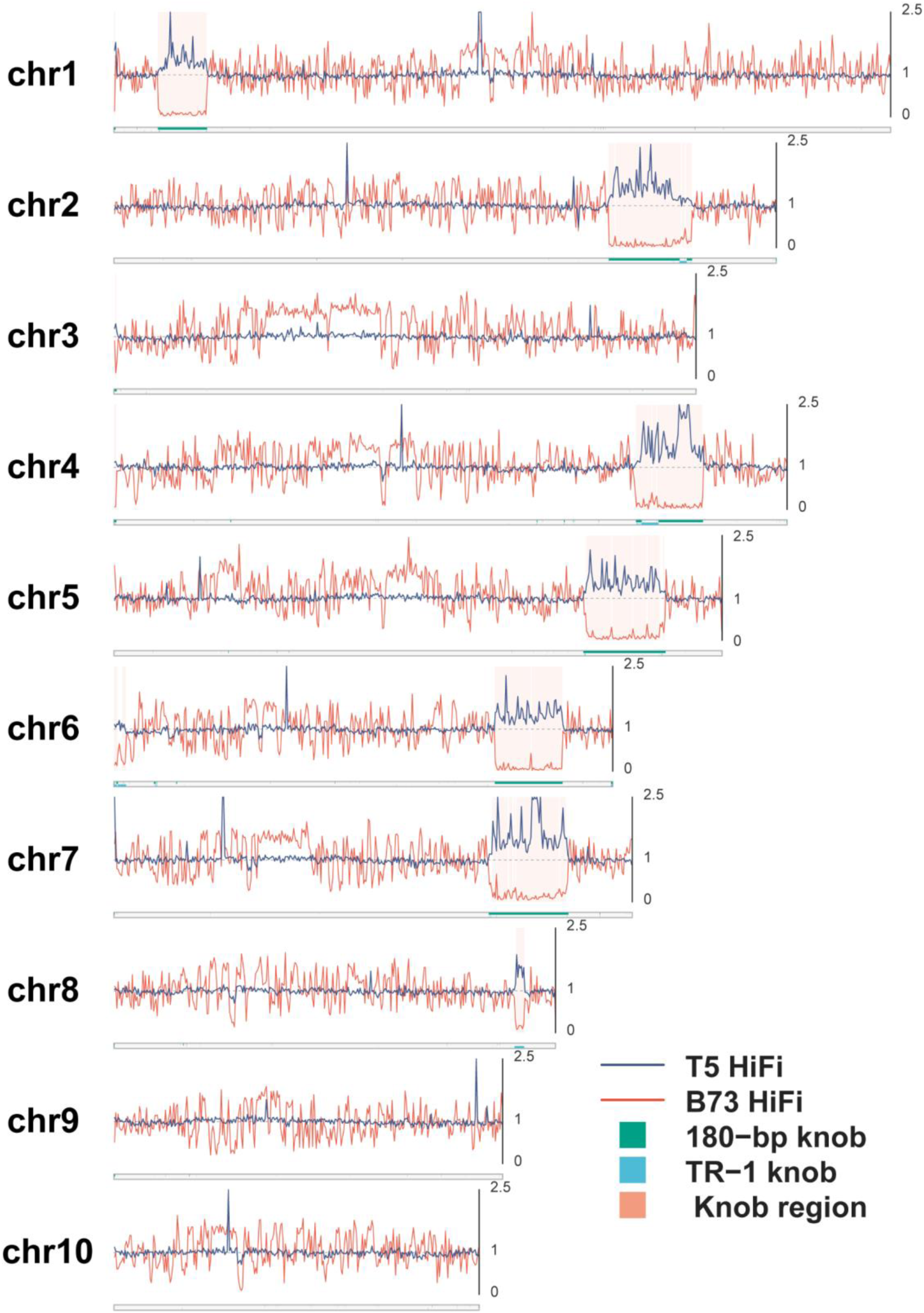
Genome-wide comparison of HiFi read coverage across knob repeat regions in Tongxi 5 (T5) and B73. HiFi read coverage was calculated in 500-kb windows and normalized to the genome-wide median coverage of each dataset. The dashed horizontal line indicates a relative coverage of 1.0. T5 and B73 HiFi coverage are shown in blue and vermilion, respectively. EDTA-annotated 180-bp knob and TR-1 repeat regions are indicated in green and cyan along the chromosome ideograms. Shaded regions denote knob-associated windows in which T5 retained substantial HiFi coverage (relative coverage ≥ 0.75) whereas B73 coverage was strongly depleted (relative coverage ≤ 0.25).

**Figure S6.**
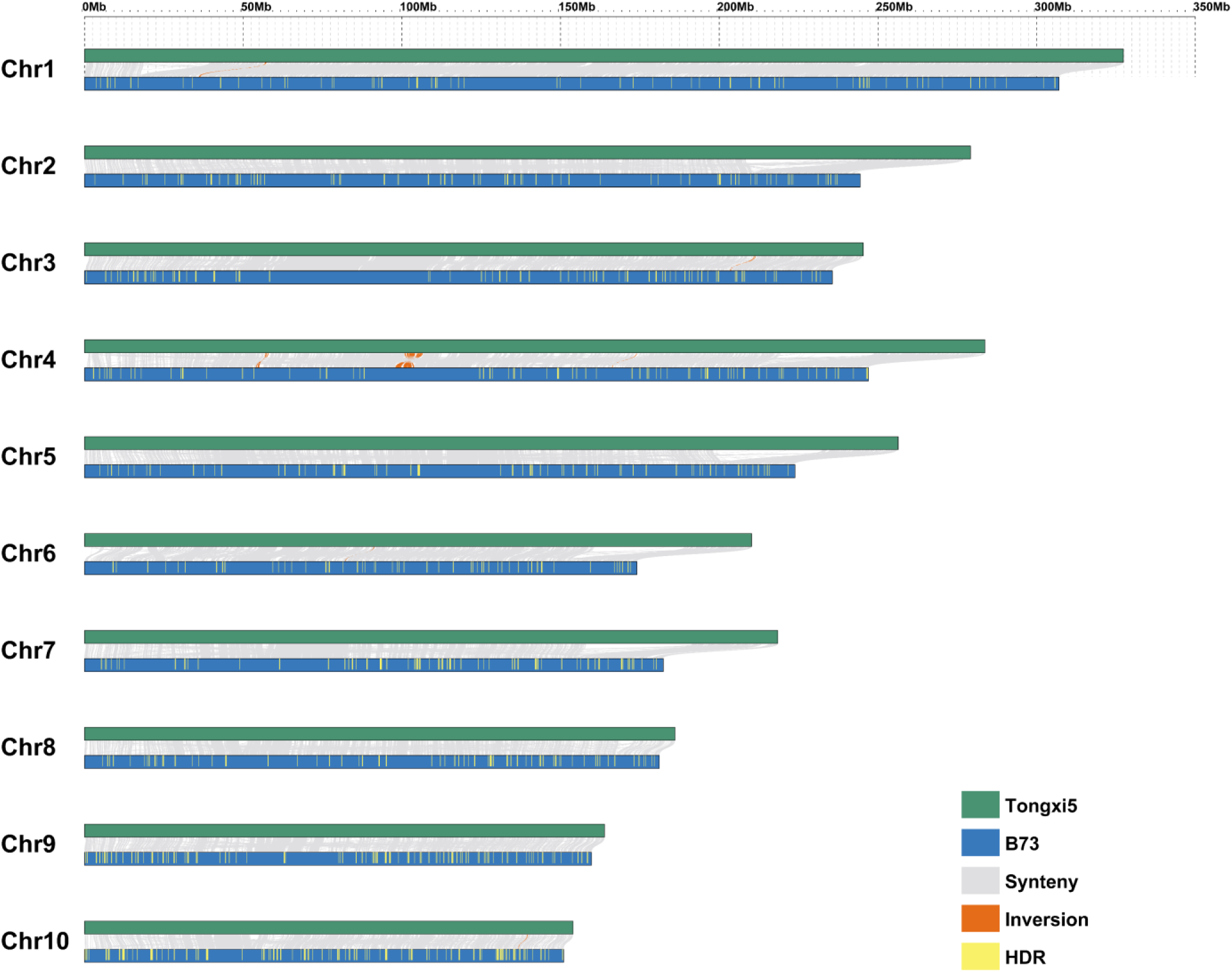
Genome-wide synteny and highly divergent regions between B73 and Tongxi5. Whole-genome pairwise alignments between B73 and Tongxi5 was visualized using GenomeSyn. For each chromosome, B73 is shown in blue and the corresponding parental genome is shown in green. Grey ribbons represent syntenic alignments between homologous chromosomal regions, whereas orange ribbons indicate inversion signals retained after filtering. Yellow vertical marks denote highly divergent regions (HDRs) larger than 100 kb on the B73 reference coordinate system. Chromosome positions are shown in megabases (Mb). The comparison highlights extensive genome-wide collinearity between B73 and parental genomes, together with several large-scale inversion signals and dispersed HDRs across chromosomes.

**Figure S7.**
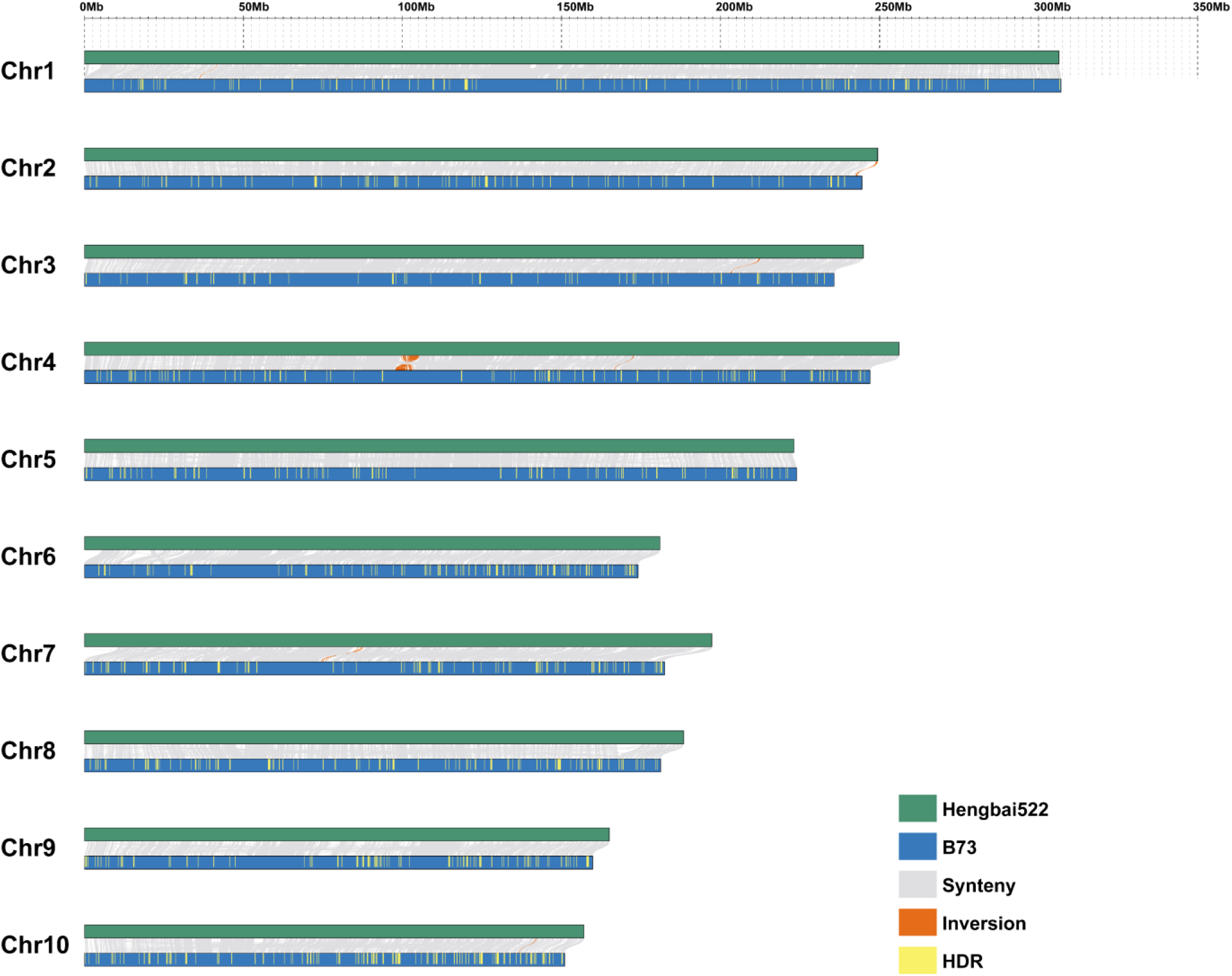
Genome-wide synteny and highly divergent regions between B73 and Hengbai522. Whole-genome pairwise alignments between B73 and Hengbai522 was visualized using GenomeSyn. For each chromosome, B73 is shown in blue and the corresponding parental genome is shown in green. Grey ribbons represent syntenic alignments between homologous chromosomal regions, whereas orange ribbons indicate inversion signals retained after filtering. Yellow vertical marks denote highly divergent regions (HDRs) larger than 100 kb on the B73 reference coordinate system. Chromosome positions are shown in megabases (Mb). The comparison highlights extensive genome-wide collinearity between B73 and parental genomes, together with several large-scale inversion signals and dispersed HDRs across chromosomes.

**Figure S8.**
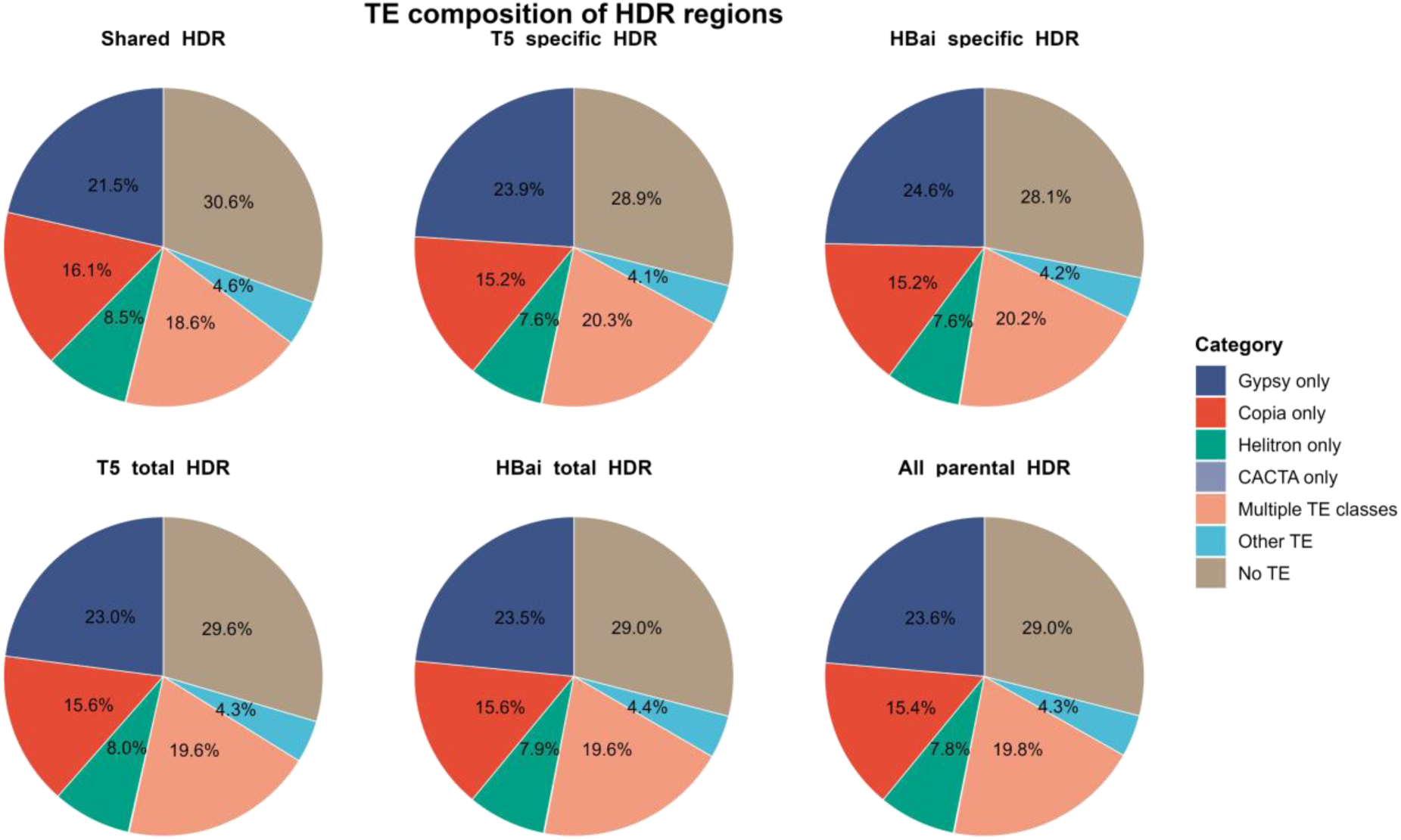
Transposable element composition of parental hyperdivergent regions. Pie charts show TE composition across shared, parent-specific and total HDRs in Tongxi 5 and Hengbai 522. Categories include Gypsy, Copia, Helitron, CACTA, multiple TE classes, other TEs and regions without TE overlap. Across all parental HDRs, 71.0% overlapped TEs, predominantly Gypsy, Copia and multi-class regions, indicating extensive association of HDRs with repeat-rich sequences. T5, Tongxi 5; HBai, Hengbai 522.

**Figure S9.**
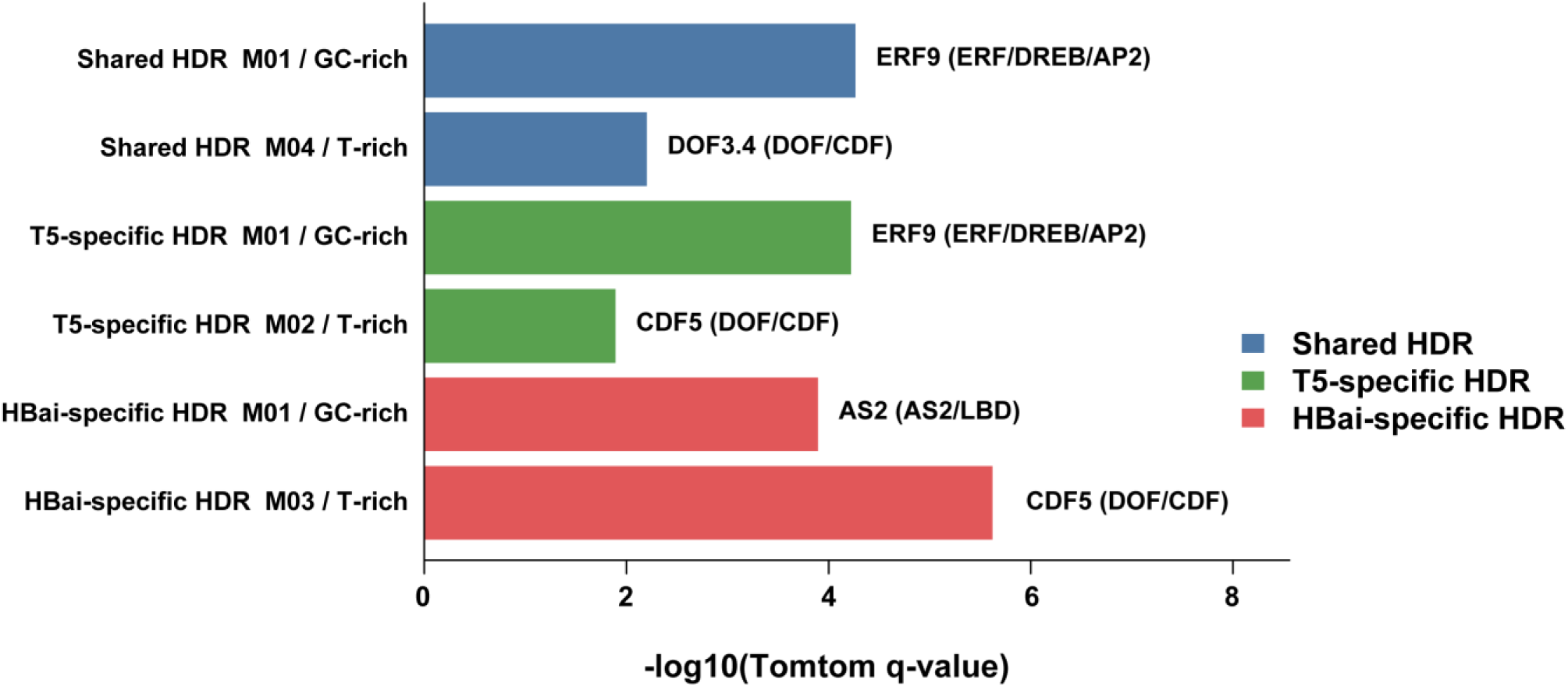
Representative transcription factor motif matches at HDR boundaries. Representative GC-rich and T-rich de novo motifs identified from 50-bp sequences flanking highly divergent region (HDR) boundaries were compared with known plant transcription factor motifs using Tomtom. Bar lengths indicate the significance of the best Tomtom match for each representative motif, expressed as −log10(q-value). Shared and T5-specific HDRs contained GC-rich motifs most similar to ERF9, an ERF/DREB-related profile, whereas HBai-specific HDRs showed a GC-rich motif most similar to AS2. T-rich motifs were mainly similar to DOF/CDF-family profiles, including DOF3.4 and CDF5. Colors indicate HDR categories: Shared HDRs, T5-specific HDRs and HBai-specific HDRs. T5, Tongxi 5; HBai, Hengbai 522.

**Figure S10.**
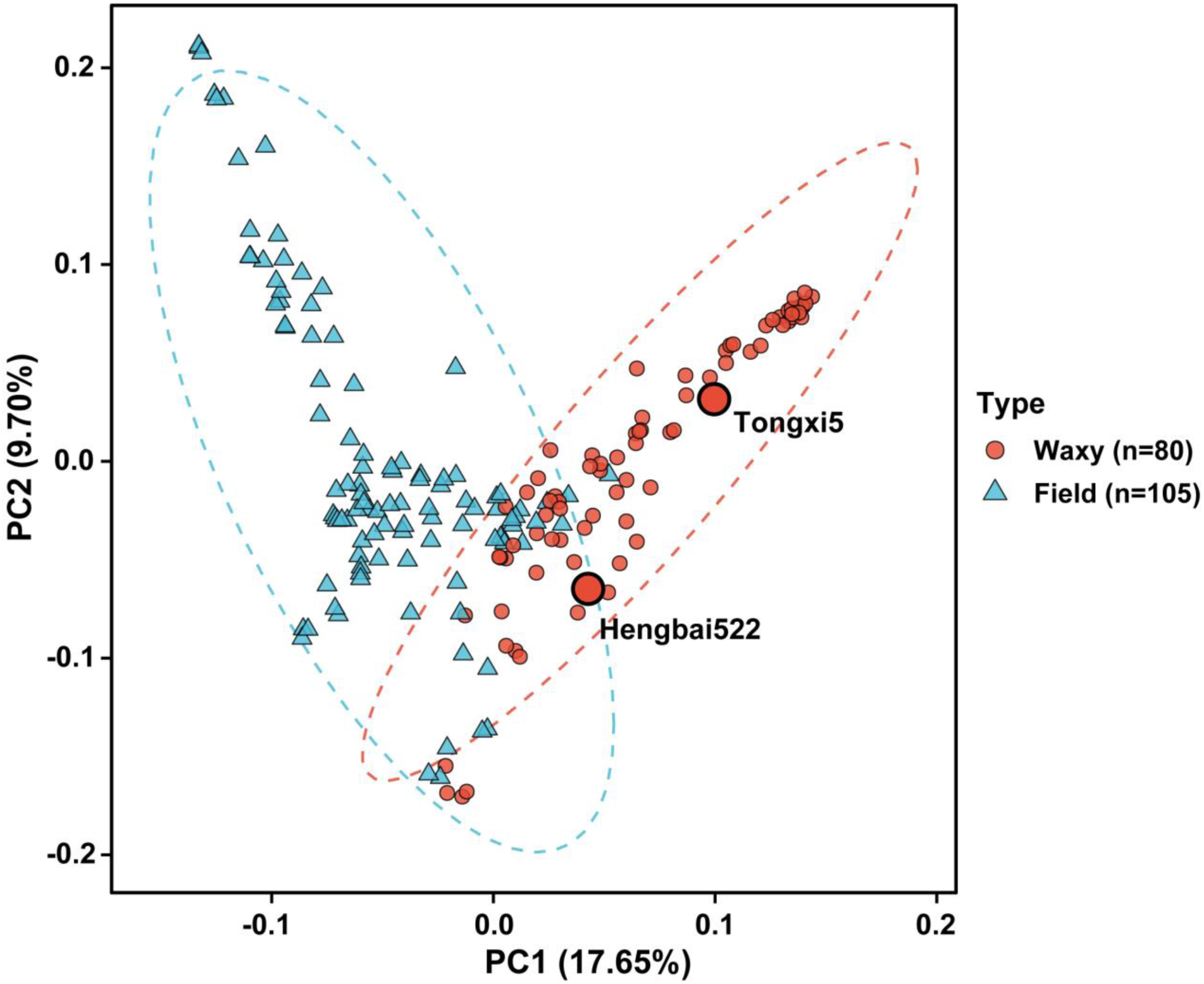
Principal component analysis of waxy and field maize inbred lines. Principal component analysis (PCA) of 185 maize inbred lines based on genome-wide SNP variation. Waxy maize (n = 80) and field maize (n = 105) are shown as red circles and cyan triangles, respectively, with dashed ellipses indicating the overall distribution of each group. Tongxi 5 and Hengbai 522 are highlighted with enlarged outlined circles. PC1 and PC2 explain 17.65% and 9.70% of the total genetic variation, respectively.

**Figure S11.**
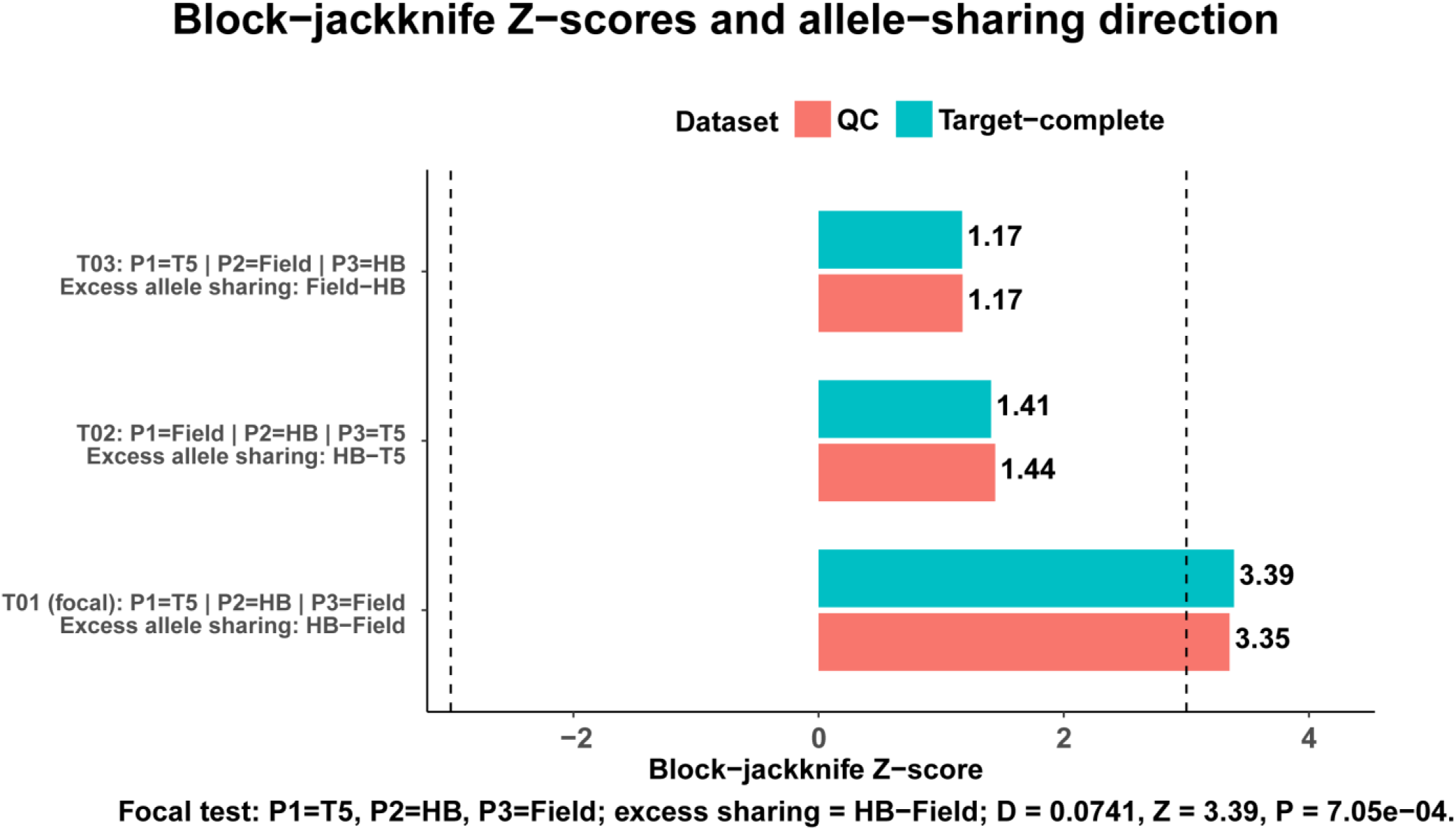
Patterson’s D-statistic tests reveal asymmetric allele sharing between the parental lines and field maize. Block-jackknife Z scores are shown for three alternative P1–P2–P3 arrangements using the quality-controlled (QC) and target-complete datasets. For each test, the corresponding P1, P2 and P3 assignments and the inferred excess allele-sharing pair are indicated. In the focal test (T01; P1 = Tongxi 5, P2 = Hengbai 522 and P3 = field maize), positive D indicates excess allele sharing between Hengbai 522 and field maize relative to Tongxi 5 and field maize. This asymmetry was significant in both datasets (Z = 3.35 for QC and Z = 3.39 for target-complete), with the target-complete dataset yielding D = 0.0741 and P = 7.05 × 10⁻⁴. Dashed lines indicate the significance threshold of ∣ Z ∣= 3.

**Figure S12.**
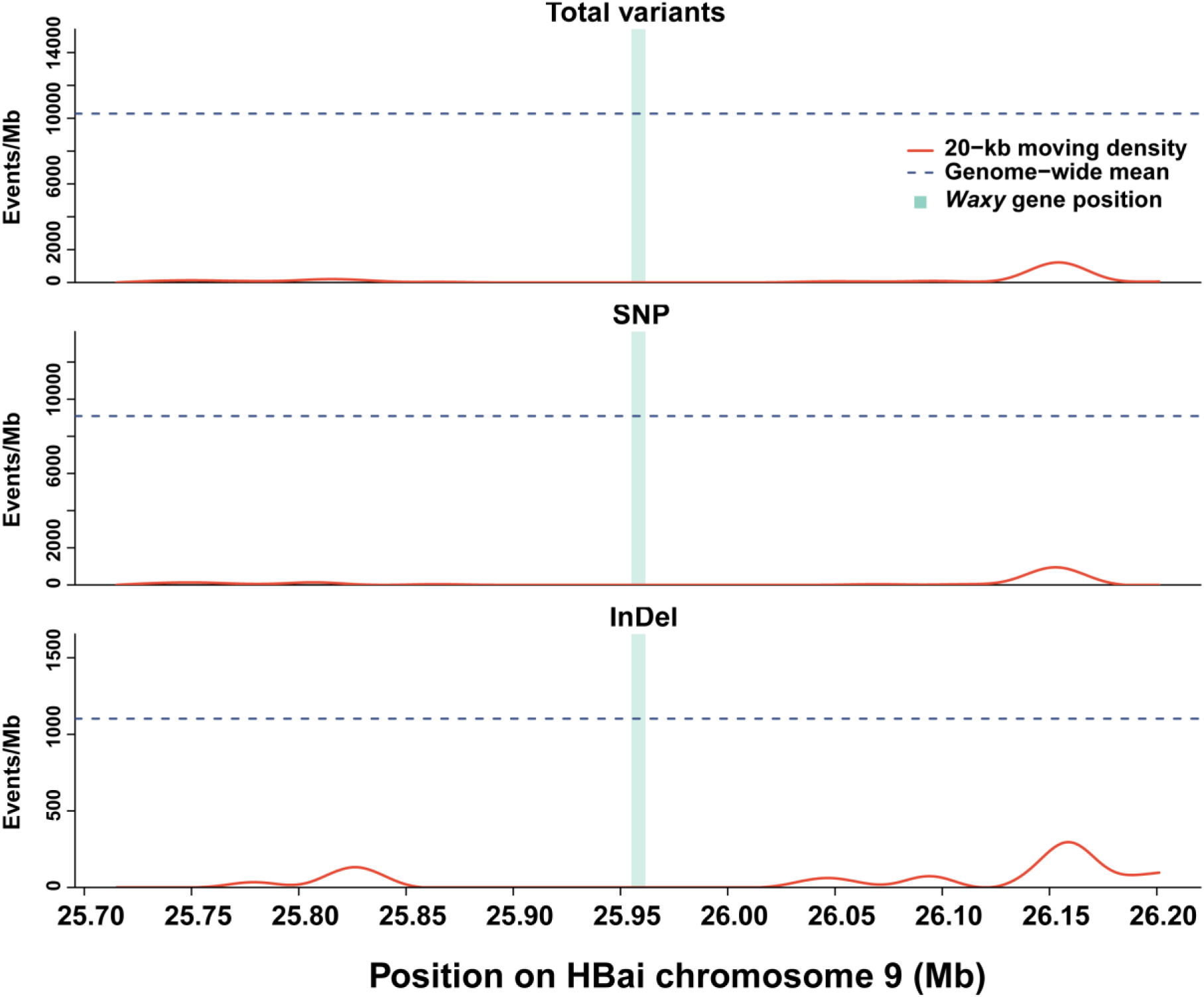
Local density of sequence variants between Hengbai 522 and Tongxi 5 around the chromosome 9 *waxy* gene region. Local densities of total variants, SNPs and InDels across HBai chromosome 9 from 25,705,078 to 26,211,605 bp. Variant density is expressed as events per Mb and was calculated using 20-kb moving windows comprising ten consecutive 2-kb bins. Red curves indicate smoothed local variant density, blue dashed lines indicate the corresponding genome-wide mean density, and the light-blue shaded region marks the *waxy* gene interval.

**Figure S13.**
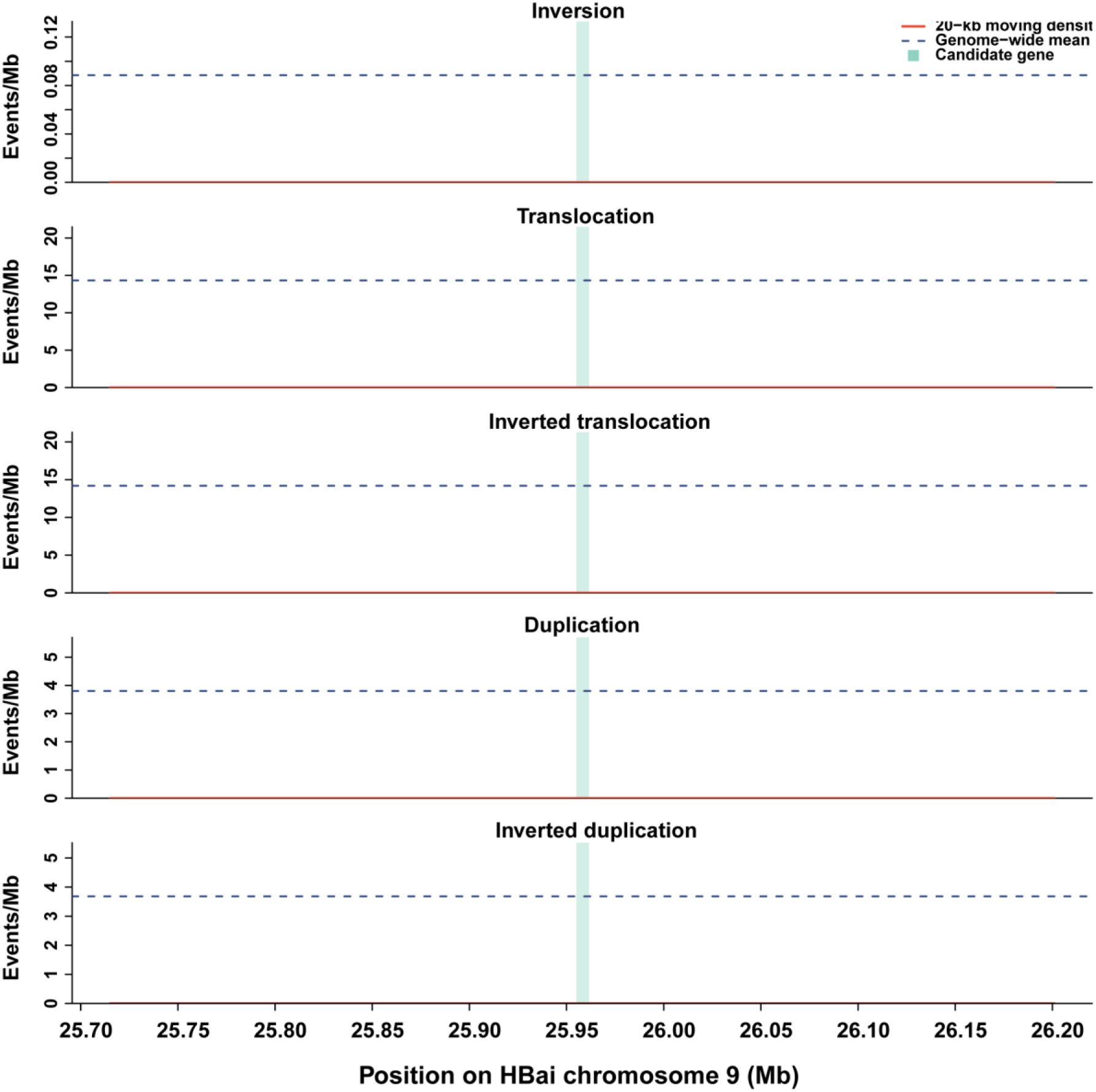
Local density of structural variations between Hengbai 522 and Tongxi 5 around the chromosome 9 *waxy* gene region. Local densities of inversions (INV), translocations (TRANS), inverted translocations (INVTR), duplications (DUP) and inverted duplications (INVDP) across HBai chromosome 9 from 25,705,078 to 26,211,605 bp. Variant density is expressed as events per Mb and was calculated using 20-kb moving windows comprising ten consecutive 2-kb bins. Red curves indicate smoothed local variant density, blue dashed lines indicate the corresponding genome-wide mean density, and the light-blue shaded region marks the *waxy* gene interval.

**Figure S14.**
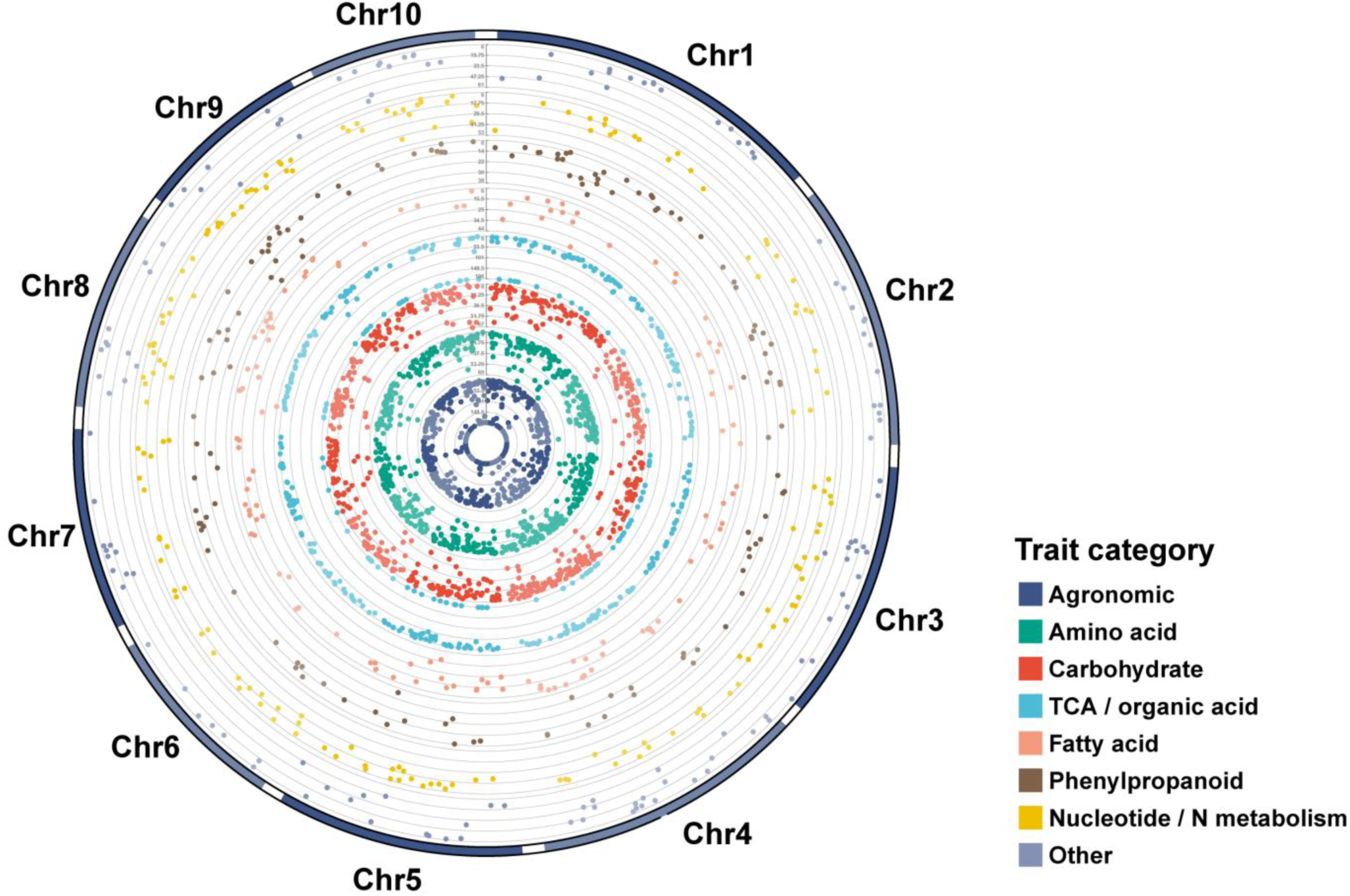
Circular Manhattan plot of significant QTLs associated with agronomic and metabolic traits. Among the 3,625 non-redundant loci, 2,886 were significant (SIG), including 2,267 SNP/InDel and 619 SV loci, whereas 739 were suggestive (SUG). Significant loci were grouped into eight agronomic and metabolic categories and visualized as concentric tracks using the CMplot R package. Within each category, loci associated with multiple traits were represented by the smallest P value; consequently, the 2,886 non-redundant loci yielded 2,916 category-level occurrences across the eight tracks. Radial height represents −log10(P), with track colors denoting trait categories and alternating shades distinguishing adjacent chromosomes.

**Figure S15.**
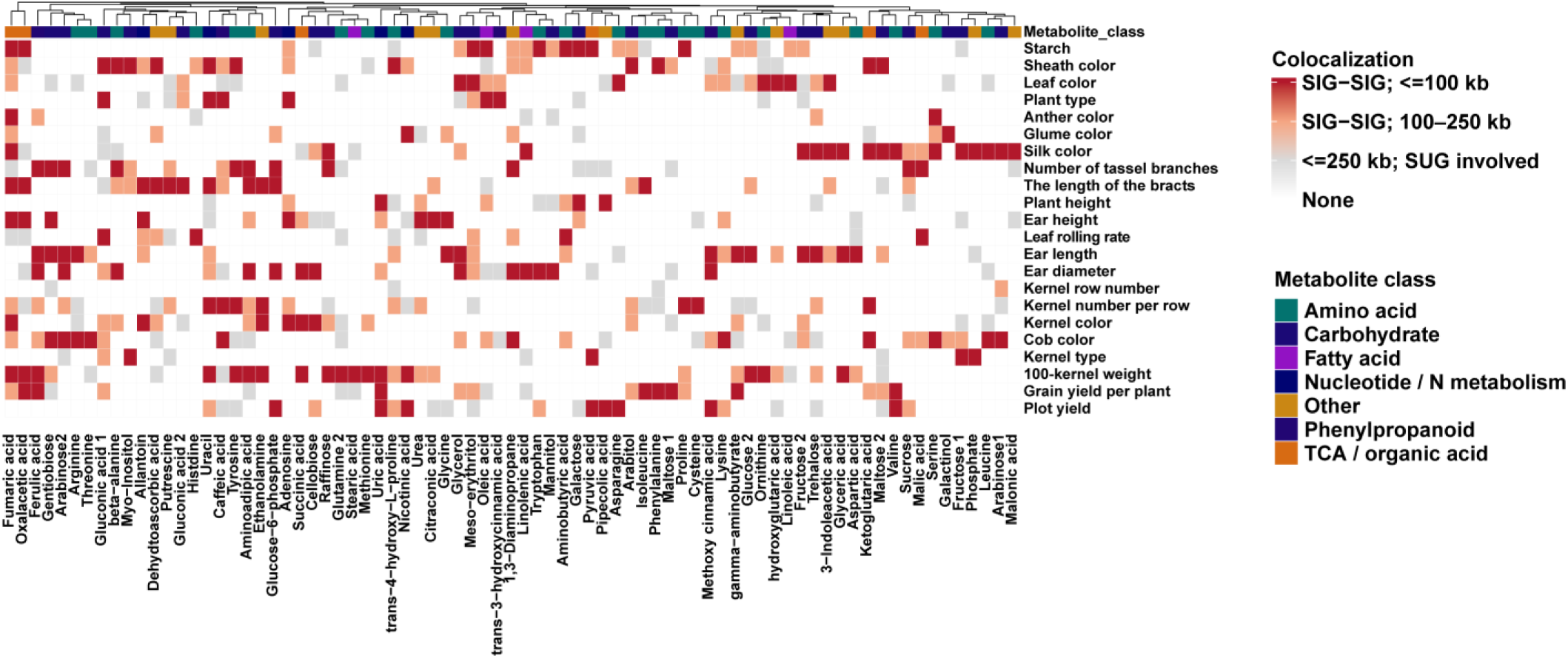
Heatmap showing agronomic–metabolic trait pairs with strong colocalization evidence. Only metabolites showing at least one significant agronomic–metabolic QTN pair within 100 kb are retained. Metabolites are annotated according to major biochemical classes, including carbohydrates, amino acids, TCA cycle/organic acids, fatty acids, phenylpropanoids, nucleotide/nitrogen-related metabolites, and other metabolites.

**Table S1. Gene Ontology enrichment analysis of the 1,259 reciprocal best-hit gene pairs identified between Tongxi 5 and Hengbai 522.**

**Table S2. KEGG pathway enrichment analysis of the 1,259 reciprocal best-hit gene pairs identified between Tongxi 5 and Hengbai 522.**

**Table S3. List of the 185 maize inbred lines, comprising 105 field maize and 80 waxy maize inbred lines, used in this study.**

**Table S4. Genome-wide genetic differenation between field and waxy maize**

**estimated using *Fst* in 10-kb sliding windows with a 5-kb step.**

**Table S5. 38 candidate introgressed regions in Hengbai 522 supported by both ABBA-derived and Dsuite analyses.**

**Table S6. Summary of 3,734 SNP/InDel-and SV-based QTN associations for agronomic and metabolite traits in this study.**

**Table S7. Parental genotype comparison of 2,886 significant SNP/InDel-and SV-based QTLs between Tongxi 5 and Hengbai 522.**

**Table S8. 245 candidate agronomic–metabolic QTL regions defined by colocalized significant QTN associations.**

## Notes

### Competing Interest Statement

The authors have declared no competing interest.

## REFERENCES

Fang, X., Liu, H., Liu, J., Song, Y., Xu, M., Jian, X., Dong, L., Zhang, Q., Xu, L., Fan, G., Wang, Z., You, Y., Feng, T., Li, W., Li, Y., Song, R. and Lin, Z. (2025) Genome assembly and population genomic analysis reveal the genetic basis of popcorn evolution. Plant Biotechnology Journal 23, 2911–2927.

Feng, L., Sun, Z., Zhang, P., Gao, J., Fu, J., Wang, Y., Yang, Y., Jia, A., Zhao, J., Li, W., Li, J., Guo, X., Wang, A., Liu, P., Liu, H.-J., Wang, H. and Chen, Z. Pedigree-resolved genome assemblies reveal the structural and functional dynamics of tropical-temperate integration in elite maize. Plant Communications.

Hufford, M.B., Seetharam, A.S., Woodhouse, M.R., Chougule, K.M., Ou, S., Liu, J., Ricci, W.A., Guo, T., Olson, A., Qiu, Y., Della Coletta, R., Tittes, S., Hudson, A.I., Marand, A.P., Wei, S., Lu, Z., Wang, B., Tello-Ruiz, M.K., Piri, R.D., Wang, N., Kim, D.w., Zeng, Y., O’Connor, C.H., Li, X., Gilbert, A.M., Baggs, E., Krasileva, K.V., Portwood, J.L., Cannon, E.K.S., Andorf, C.M., Manchanda, N., Snodgrass, S.J., Hufnagel, D.E., Jiang, Q., Pedersen, S., Syring, M.L., Kudrna, D.A., Llaca, V., Fengler, K., Schmitz, R.J., Ross-Ibarra, J., Yu, J., Gent, J.I., Hirsch, C.N., Ware, D. and Dawe, R.K. (2021) De novo assembly, annotation, and comparative analysis of 26 diverse maize genomes. Science 373, 655–662.

Igolkina, A.A., Vorbrugg, S., Rabanal, F.A., Liu, H.-J., Ashkenazy, H., Kornienko, A.E., Fitz, J., Collenberg, M., Kubica, C., Mollá Morales, A., Jaegle, B., Wrightsman, T., Voloshin, V., Bezlepsky, A.D., Llaca, V., Nizhynska, V., Reichardt, I., Bezrukov, I., Lanz, C., Bemm, F., Flood, P.J., Nemomissa, S., Hancock, A., Guo, Y.-L., Kersey, P., Weigel, D. and Nordborg, M. (2025) A comparison of 27 Arabidopsis thaliana genomes and the path toward an unbiased characterization of genetic polymorphism. Nature Genetics 57, 2289–2301.

Li, C., Li, Z., Lu, B., Shi, Y., Xiao, S., Dong, H., Zhang, R., Liu, H., Jiao, Y., Xu, L., Su, A., Wang, X., Zhao, Y., Wang, S., Fan, Y., Luo, M., Xi, S., Yu, A., Wang, F., Ge, J., Tian, H., Yi, H., Lv, Y., Li, H., Wang, R., Song, W. and Zhao, J. (2025a) Large-scale metabolomic landscape of edible maize reveals convergent changes in metabolite differentiation and facilitates its breeding improvement. Molecular Plant 18, 619–638.

Li, K., Yu, Y., Yan, S., Li, W., Xu, J., Li, G., Li, W., Liu, J., Qi, X., Huang, W., Zhang, Q., Kong, Q., Xiao, Y., Zhang, N., Luo, J., Chen, L., Feng, L., Zhu, W., Wen, T., Xie, L., Li, Y., Lu, W., Li, C., Gui, S., Xiao, Y., Yang, N., Zhuo, L., Fernie, A.R., Liu, H.-J., Hu, J. and Yan, J. (2025b) Genetic basis of flavor complexity in sweet corn. Nature Genetics 57, 2842–2851.

Lin, Z., Qin, P., Zhang, X., Fu, C., Deng, H., Fu, X., Huang, Z., Jiang, S., Li, C., Tang, X., Wang, X., He, G., Yang, Y., He, H. and Deng, X.W. (2020) Divergent selection and genetic introgression shape the genome landscape of heterosis in hybrid rice. Proceedings of the National Academy of Sciences 117, 4623–4631.

Luo, J., He, C., Yan, S., Jiang, C., Chen, A., Li, K., Zhu, Y., Gui, S., Yang, N., Xiao, Y., Wu, S., Zhang, F., Liu, T., Wang, J., Huang, W., Yang, Y., Wang, H., Yang, W., Li, W., Zhuo, L., Fernie, A.R., Zhan, J., Wang, L. and Yan, J. (2024) A metabolic roadmap of waxy corn flavor. Molecular Plant 17, 1883–1898.

Tan, K., Liu, X., Wang, Z., Zhang, Z., Huang, W., Liu, S., Lin, Z., Zhao, H., Zhao, H., Liu, Y., Han, F., Lai, J., Song, W., Zhao, J. and Chen, J. (2026) Near-complete genome assembly of a transformation-efficient elite inbred line LH244 and its comparison with B73. Journal of Integrative Plant Biology 68, 366–382.

