## Supplementary material for "Parental Genome Assemblies of Suyunuo1 Reveal Structural Variation Underlying Edible Waxy Maize Evolution, Superior Hybrid Performance and Yield - Flavor Balance": Method-sup

**Supplemental Methods**

**Plant material, DNA library construction, and sequencing**

Tongxi 5 and Hengbai 522 seeds were obtained from the Comprehensive Germplasm Gene Bank of Jiangsu Agricultural Genetic Resources. Fresh leaves were collected at the seedling stage for genome sequencing. High-molecular-weight genomic DNA was extracted using the CTAB method (Porebski et al., 1997) and assessed by agarose gel electrophoresis, NanoDrop spectrophotometry and Qubit fluorometry. For short-read sequencing, approximately 300-bp paired-end libraries were constructed using the VAHTS Universal Plus DNA Library Prep Kit for MGI V2 following Covaris fragmentation. For Oxford Nanopore ultra-long sequencing, libraries were prepared using the SQK-LSK110 ligation sequencing kit according to the manufacturer’s protocol, loaded onto R9.4.1 Spot-On Flow Cells and sequenced on a PromethION platform (Oxford Nanopore Technologies, Oxford, UK) for 72 h. For PacBio HiFi sequencing, SMRTbell libraries with an insert size of approximately 15 kb were prepared using the SMRTbell Express Template Prep Kit 2.0 (Pacific Biosciences, USA) and sequenced on a PacBio Sequel II platform with 30-h runs. Oxford Nanopore and PacBio sequencing were performed at Wuhan Benagen Technology Co., Ltd. (Wuhan, China).

**Plant material, RNA library construction, sequencing**

Total RNA was extracted from multiple tissues of Tongxi 5 and Hengbai 522, including roots, embryos, endosperm, buds, pollen, ear axes, and leaves, using the RNAprep Pure Plant Plus Kit (Tiangen Biotech, Beijing, China) according to the manufacturer’s instructions. RNA-seq libraries were constructed using the NEBNext Ultra RNA Library Prep Kit for Illumina (NEB, USA) and sequenced on an Illumina platform to generate 150-bp paired-end reads at Wuhan Benagen Technology Co., Ltd., Wuhan, China.

**Hi-C sequencing and data processing**

High-quality genomic DNA from young leaves of Tongxi 5 and Hengbai 522 was used for Hi-C sequencing. Chromatin was cross-linked with formaldehyde, and in situ Hi-C libraries were constructed following the DNase-based protocol described by Ramani et al (Ramani et al., 2020). The libraries were sequenced on an Illumina NovaSeq platform to generate 150-bp paired-end reads.

**Gap-free genome de novo assembly**

ONT ultra-long reads were independently assembled using NextDenovo v2.5.2 with the parameters read_cutoff = 1k, sort_options = -m 9g -t 6 -k 40, and minimap2_options_cns = -t 21 -k17 -w17 (Hu et al., 2024). PacBio reads were processed using CCS with default settings to generate circular consensus sequencing reads (HiFi reads). HiFi-only and hybrid assemblies integrating PacBio HiFi and ONT ultra-long reads were subsequently generated using hifiasm (Cheng et al., 2021). Residual haplotypic duplications in the selected assembly were identified and removed using purge_dups v1.2.5 based on read-depth and sequence-alignment evidence (Guan et al., 2020). Potential contaminant contigs were identified by sequence alignment using minimap2 (Li, 2018) and removed according to sequence identity and alignment coverage. Hi-C reads were processed and mapped to the assembled contigs using Juicer v1.6 (Durand et al., 2016b). Valid Hi-C contacts were used for misassembly correction and chromosome-scale clustering, ordering and orientation with 3D-DNA v180419 (Dudchenko et al., 2017). The resulting contact maps were manually inspected using Juicebox v1.11.08, and contig order and orientation were corrected where necessary (Durand et al., 2016a). The manually curated contig order and orientation were then incorporated to generate chromosome-level assemblies. Residual assembly gaps were subsequently filled using a hierarchical strategy based on alternative corrected genome assemblies, ONT sequences and PacBio HiFi reads. Alternative assemblies generated by NextDenovo, hifiasm or other assembly approaches were given priority, followed by ONT consensus or raw reads and then HiFi CCS reads. Candidate gap-filling sequences were aligned to the chromosome-level assemblies using Winnowmap (Jain et al., 2020), and alignments spanning both flanks of each gap were identified. For each gap, the longest high-confidence alignment completely bridging the gap was selected to replace the corresponding gap-containing sequence. Gap-filled intervals were further validated by remapping alternative assemblies, ONT reads and HiFi reads. A replacement was considered supported when independent sequence evidence spanned both boundaries of the filled interval, confirming sequence continuity across the newly incorporated region. Only supported replacements were retained in the gap-free assemblies. The gap-filled assemblies were subsequently polished with PacBio HiFi reads using Racon v1.4.13 (https://github.com/lbcb-sci/racon).

ONT ultra-long reads were aligned to the genome using Winnowmap (Jain et al., 2020), and uniquely mapped reads overlapping the terminal 50 bp of each chromosome were extracted. Telomeric repeat motifs (CCCTAAA/TTTAGGG) were counted in these reads, and the read containing the highest number of repeats was selected as the reference for local consensus generation with medaka_consensus (https://github.com/nanoporetech/medaka). The resulting consensus sequences were aligned to the corresponding chromosome ends using mucmer (Marçais et al., 2018). Based on alignment identity, coverage, coordinates, and the continuity of telomeric repeat arrays, supported terminal sequences were incorporated into the assembly while minimizing changes to the original chromosome sequence. The supported terminal sequences were incorporated to refine chromosome ends, generating the final gap-free chromosome-level assemblies. Hi-C reads were remapped to the final assemblies, and genome-wide contact matrices were constructed at a resolution of 150 kb using HiCExplorer v3.6 (Wolff et al., 2020). The interaction matrices were subsequently visualized as Hi-C contact heatmaps.

**Assembly quality assessment**

Genome assembly quality was evaluated in terms of completeness, contiguity, and sequence accuracy. Gene-space completeness was assessed using BUSCO v5.4.7 in genome mode (Manni et al., 2021). Assembly statistics, including contig and scaffold N50 values, were calculated using QUAST (Gurevich et al., 2013). The continuity of repeat-rich genomic regions was further evaluated using the long terminal repeat assembly index (LAI) (Ou et al., 2018). Consensus quality values and k-mer completeness were estimated using Merqury v1.3 based on k-mers derived from the sequencing reads (Rhie et al., 2020).

**Repetitive sequence annotation**

RepeatModeler (v2.0.6) (Flynn et al., 2020) was used to identify repetitive elements directly from the assembled genome. Long terminal repeat retrotransposons were independently predicted using LTR_FINDER (Xu and Wang, 2007) and LTRharvest (v1.6.5) (Ellinghaus et al., 2008), and the resulting candidates were integrated and filtered using LTR_retriever (v3.0.1) (Ou and Jiang, 2018) to generate a nonredundant LTR library. The RepeatModeler and LTR libraries were subsequently merged, and unclassified elements were further classified using TEclass (Abrusán et al., 2009). Transposable elements were identified using RepeatMasker with the combined de novo and RepBase libraries, supplemented by protein homology-based annotation using RepeatProteinMask (Tempel, 2012). All annotations were subsequently merged and deduplicated to generate the final set of interspersed repeats. Knob repeats in Tongxi 5, Hengbai 522 and 26 field maize genomes were further annotated using the Extensive de novo TE Annotator (EDTA) with maize-specific settings and a curated maize repeat library (Ou et al., 2019). For Pacbio long reads alignment, canonical knob-associated repeats were identified from the EDTA annotations as the 180-bp knob repeat (knob180) and TR-1 repeat families, and their genomic coordinates were extracted separately to characterize repeat abundance and chromosomal distribution. HiFi read coverage from Tongxi 5 and B73 was summarized in matched 500-kb windows and normalized to the genome-wide median coverage of each dataset. Knob180 and TR-1 annotations were overlaid with the normalized coverage profiles, and knob-associated windows with relative coverage ≥0.75 in Tongxi 5 and ≤0.25 in B73 were defined as knob regions. Knob annotations were further intersected with Tongxi 5-specific insertions to assess knob-repeat enrichment relative to the corresponding regions in Hengbai 522 and B73. For cytological validation, mitotic metaphase chromosome spreads of Tongxi 5, Hengbai 522, and B73 were prepared with minor modifications from previous studies (Du et al., 2016; Kato, 1999; Zhu et al., 2017). Six oligonucleotide probes derived from tandem repeat sequences, namely Gypsy-2, MR68-3, Knob-2, (ACT)10, 5S-1, and 5S-2, were utilized for fluorescence in situ hybridization (FISH) to distinguish individual maize chromosomes according to previously established procedures (Chen et al., 2019; Zhu et al., 2017). Microscopic images were captured using an Olympus BX53 fluorescence microscope equipped with SPOT CCD camera (SPOT Cooled Color Digital, DP72, Olympus, Japan).

**Non-coding RNA annotation**

Transfer RNA genes were predicted using tRNAscan-SE (v 2.0.12) (Lowe and Eddy, 1997) based on their conserved structural features. Ribosomal RNA genes were identified using RNAmmer (Lagesen et al., 2007). Other non-coding RNAs, including small nuclear RNAs and microRNAs, were annotated using INFERNAL (v 1.1.4) (Nawrocki and Eddy, 2013) by searching against the Rfam database.

**Gene annotation**

Protein-coding genes were annotated by integrating transcriptome-based, protein homology-based, and ab initio predictions. For transcriptome-based annotation, Illumina RNA-seq reads were quality-filtered using fastp (Chen et al., 2018) and aligned to the repeat-masked genome using HISAT2 (v2.2.1) (Kim et al., 2019). The resulting alignments were assembled into transcript models using StringTie (v2.2.1) (Pertea et al., 2015), and candidate coding regions were identified using TransDecoder (v5.7.0) (<https://github.com/TransDecoder/TransDecoder>). For homology-based prediction, protein sequences from closely related plant species and conserved proteins from the BUSCO database were aligned to the genome using miniprot (v0.13) (Li, 2023). Ab initio gene prediction was performed on the repeat-masked genome using AUGUSTUS (v3.5.0) (Stanke et al., 2008) and GlimmerHMM (v3.0.4) (Delcher et al., 2007), with the prediction models trained or optimized using transcriptomic and protein homology evidence. Transcript-derived gene models, protein homology alignments, and ab initio predictions were integrated using EVidenceModeler (v2.1.0) (Haas et al., 2008) to generate a consensus gene set. The resulting gene models were further filtered based on coding-sequence integrity, the presence of valid start and stop codons, gene length, and supporting evidence. Gene models supported by multiple evidence sources were preferentially retained to generate the final high-confidence protein-coding gene annotation.

**Transcriptome-based evaluation of waxy maize reference genomes**

Publicly available paired-end RNA-seq datasets from two independent waxy maize studies, comprising nine Jingkenuo 2000 libraries (PRJNA1050395) (Zhao et al., 2025) and twelve SWL01 libraries (PRJNA625943) (Gu et al., 2020), were downloaded from the European Nucleotide Archive. Raw reads were quality-filtered using fastp v0.23.4 (Chen et al., 2018) and independently aligned to the B73 RefGen_v4, Tongxi 5 and Hengbai 522 genomes using STAR v2.7.11b (Dobin et al., 2013). STAR genome indices were constructed using the corresponding GTF annotations with --sjdbOverhang 149, and all libraries were mapped using identical parameters with 12 threads. Mapping performance was analyzed separately for Jingkenuo 2000 and SWL01, with uniquely mapped read proportion used as the primary measure of reference compatibility. Paired differences among the three reference genomes were evaluated using Wilcoxon signed-rank tests.

**Identification of genetic variants and syntenic analysis**

Pairwise whole-genome alignments among the B73 RefGen_v4, Hengbai 522, and Tongxi 5 genomes were performed chromosome by chromosome using NUCmer from the MUMmer package (Marçais et al., 2018). The resulting alignments were filtered using delta-filter with parameters “-m -i 90 -l 100” to retain high-confidence alignments with ≥90% sequence identity and a minimum alignment length of 100 bp. Structural variations were subsequently identified using SyRI (v1.6.3) (Goel et al., 2019) with default settings and classified into insertions, deletions, duplications, inversions, translocations, and other genomic rearrangements. Syntenic regions were defined according to SyRI SYN annotations, and their non-overlapping genomic intervals were used to quantify syntenic coverage in each pairwise genome comparison.

**Homology inference among protein-coding genes of Tongxi 5, Hengbai 522 and 26 field maize genomes**

Protein-coding genes from Tongxi 5, Hengbai 522 and 26 field maize reference genomes were analyzed using GeneTribe v1.2.1 (Chen et al., 2020). Protein sequences, gene coordinates and chromosome information were formatted as FASTA, BED and chromosome-list files, respectively, with gene identifiers standardized across genomes. Tongxi 5 and Hengbai 522 were independently compared with each of the 26 field maize genomes using genetribe core, which integrates sequence similarity and genomic collinearity for homolog inference. Genes consistently classified as singletons across all 26 comparisons were considered to lack detectable homologs in field maize. These genes were further classified using the Tongxi 5–Hengbai 522 comparison: genes remaining singletons between the two parental genomes were designated parental-specific candidates, whereas homologous pairs whose members both lacked detectable homologs in all 26 field maize genomes were defined as shared-specific candidates. Among the latter, reciprocal best-hit (RBH) pairs, in which each gene was the best homologous match of its counterpart in the other parental genome, were retained as a high-confidence set of shared-specific homologs. GO and KEGG annotations were obtained from eggNOG-mapper annotations (Huerta-Cepas et al., 2017) generated for the complete protein-coding gene sets of Tongxi 5 and Hengbai 522. GO and KEGG pathway enrichment analyses were performed separately for parental-specific genes and for the Tongxi 5 and Hengbai 522 members of the 1,259 shared-specific RBH pairs, using the corresponding genome-wide annotated gene sets as backgrounds. For the shared-specific genes, terms or pathways significantly enriched in both parental genomes were considered common enrichments. Enrichment analyses were conducted using the enricher function in the R package clusterProfiler (Xu et al., 2024). *P-*values were adjusted for multiple testing using the Benjamini–Hochberg method, and GO terms and KEGG pathways with an adjusted *P* value (FDR) < 0.05 were considered significantly enriched.

**Identification and annotation of hyperdivergent regions**

Pairwise genome comparisons were performed between B73 and each parental line, Tongxi 5 and Hengbai 522, using B73 as the common coordinate reference. Hyperdivergent regions (HDRs) were extracted from SyRI outputs (Goel et al., 2019), retaining only those nested within syntenic alignment blocks to ensure positional orthology. Visualization of chromosomal distributions and length compositions was performed via the GenomeSyn package (Zhou et al., 2022). To quantify transposable-element (TE) content, B73 TE annotations were converted from GFF3 to BED format and intersected with shared and parental-line-specific syntenic HDRs using BEDTools (Quinlan and Hall, 2010). TE identities and classes were summarized for each HDR category. To account for overlapping and nested annotations, TE intervals were merged into non-redundant regions, and their cumulative coverage within each HDR category was calculated from overlap lengths. To investigate sequence features associated with HDR boundaries, HDRs were classified into shared HDRs, Tongxi 5-specific HDRs and Hengbai 522-specific HDRs. For each HDR category, 50-bp flanking sequences immediately adjacent to both sides of each HDR were extracted from the B73 reference genome using BEDTools (Quinlan and Hall, 2010). De novo motif discovery was performed separately for each HDR category using MEME in the MEME Suite v5.5.9 with DNA mode and the zero-or-one-occurrence-per-sequence model (Bailey et al., 2006). Motifs with MEME E-values < 0.05 were retained as significant boundary-associated motifs. To infer potential transcription factor binding similarities, significant motifs were further compared with the JASPAR 2026 CORE plant non-redundant motif database using Tomtom (Rauluseviciute et al., 2024), and matches with q-values < 0.05 were considered significant.

**Whole-genome short-read resequencing and variant calling**

Genomic resequencing data from 185 maize inbred lines, including 105 field maize and 80 waxy maize accessions, were analyzed using B73 RefGen_v4 as the reference genome. Quality-filtered paired-end reads were aligned to the reference using BWA-MEM2 (v2.2.1) (Li and Durbin, 2009) with default parameters. Alignments were converted to BAM format, sorted and indexed using SAMtools (v1.17) (Li et al., 2009), and PCR duplicates were subsequently removed. SNPs and InDels were called for each accession using GATK HaplotypeCaller (v4.5.0.0) in GVCF mode (McKenna et al.), followed by joint genotyping across all samples. Variants were filtered using the criteria QD < 2.0, MQ < 40.0, FS > 60.0, SOR > 3.0, MQRankSum < −12.5 or ReadPosRankSum < −8.0. Sites with a minor allele frequency (MAF) < 0.05 or genotype missing rate > 0.1 were further excluded. The final high-quality dataset contained 6,879,860 SNPs and 467,232 InDels and was used for subsequent population genomic analyses.

**Population genomic and introgression analyses**

Principal component analysis (PCA) was performed using GCTA (Yang et al., 2011). Briefly, the genotype matrix was used to construct a genomic relationship matrix using the --make-grm function, and principal components were subsequently obtained by eigenvalue decomposition of the relationship matrix using --pca. The first two principal components were plotted to characterize population structure and genetic differentiation between field and waxy maize and to determine the relative genetic relationships of Tongxi 5 and Hengbai 522 with the two germplasm groups. Population differentiation between field and waxy maize was quantified using the Weir–Cockerham *F_ST_* estimator implemented in VCFtools (Danecek et al., 2011), and the genome-wide mean *F_ST_* was calculated across polymorphic sites.

To test for asymmetric allele sharing between the two parental lines and field maize, Patterson’s D statistic was calculated using Dsuite (Malinsky et al., 2021). The analysis included 20 accessions comprising Tongxi 5, Hengbai 522, ten field maize inbred lines and eight teosinte accessions used as the outgroup. For the focal test, Tongxi 5, Hengbai 522 and field maize were assigned as P1, P2 and P3, respectively, corresponding to the topology (((Tongxi 5, Hengbai 522), field maize), outgroup). Statistical significance was assessed using block jackknifing, with the resulting Z score and associated two-sided *P* value used to evaluate deviation from *D* = 0. Local patterns of allele sharing were subsequently examined using two complementary window-based approaches. First, Dsuite Dinvestigate (Malinsky et al., 2021) was applied using windows containing 1,000 informative SNPs with a step of 500 SNPs to calculate local statistics. Second, ABBABABAwindows.py (Martin et al., 2015) was used to perform a fixed-coordinate genome scan with 1-Mb windows and a 500-kb step. Sites were required to satisfy a minimum data proportion of 0.80 within each population, and windows containing fewer than 100 informative biallelic SNPs were excluded. Candidate Hengbai 522–field maize shared regions were identified using the empirical genome-wide distribution of *f_dm*. Windows were first required to have positive D and *f_dm* values and an *f_dm* value within the upper 5% of the genome-wide distribution. To minimize the influence of isolated extreme windows and local stochastic variation, candidate regions were further required to contain at least three consecutive overlapping outlier windows. Candidate regions independently identified by the two approaches were compared based on their genomic coordinates, and regions showing concordant positive signals were retained. Overlapping supported intervals were merged to define the final set of candidate introgressed regions. Genes overlapping these regions were extracted from the corresponding maize genome annotation and assessed for previously characterized functions.

**Sequence variation surrounding the *waxy* locus**

Genome-wide sequence variation between Tongxi 5 and Hengbai 522 was characterized based on their pairwise whole-genome alignment. All genetic variants identified from the genome comparison using SyRI were summarized in non-overlapping 2-kb windows across the genome. To assess sequence conservation surrounding the *waxy* locus, variant density within a 300-kb interval encompassing the *waxy* gene was compared with the genome-wide background. Differences in variant density between the *waxy* interval and the genomic background were evaluated using Welch’s two-sample t-test. The reduction in variant density was calculated as 100 × (1 − mean variant density within the waxy region/mean genome-wide variant density).

**Agronomic performance investigation and GC–MS metabolite profiling**

Maize inbred lines were grown in the Liuhe Base of Jiangsu Academy of Agricultural Sciences (32°29′ N, 118°50′ E) in 2025 using a randomized complete block design. Twenty-two agronomic related traits were evaluated in the field at the appropriate developmental stages, including plant height, ear height, plant type, number of tassel branch, the length of the bracts, ear length, ear diameter, kernel row number, kernel number per row, kernel type, 100-kernel weight, grain yield per plant, plot yield, seed starch, glume color, silk color, kernel color, cob color, leaf color, sheath color, anther color, leaf rolling rate. For kernel metabolite profiling, approximately twenty uniform kernels were collected from the middle portion of each ear at 22 days after pollination. Kernel samples were freeze‑dried under liquid nitrogen prior to metabolite extraction. For GC–MS analysis, 50 mg of lyophilized kernel powder was extracted according to a previously established protocol with minor modifications (Luo et al., 2024). The dried extracts were derivatized with N-methyl-N-(trimethylsilyl)trifluoroacetamide as described previously (Yan et al., 2018), and analyzed using an Agilent 7890A gas chromatograph coupled to a 5975C mass spectrometer (Agilent Technologies, USA). One microliter of each derivatized sample was injected at 270 °C in split mode (50:1), with high-purity helium (>99.999%) as the carrier gas at a constant flow rate of 1 mL min⁻¹. Metabolites were separated on a DB-35MS UI capillary column (30 m × 0.25 mm, 0.25 μm). GC–MS data processing and metabolite quantification were performed following the procedure described by Yan et al. (2018) (Yan et al., 2018) using Agilent MassHunter Quantitative Analysis software (version B.07.01).

**Structural variant genotyping and functional annotation across 185 maize inbred lines**

A non-redundant set of structural variants was used to genotype SVs across the 185 maize inbred lines. To reduce reference-mapping bias around structurally divergent regions, the B73 RefGen_v4 sequence and the merged SV dataset were incorporated into a graph-based reference using vg (Hickey et al., 2020). The quality-filtered paired-end reads from each accession were mapped independently to the graph using the Giraffe workflow (Sirén et al.), and read support for individual SV alleles was summarized from the resulting graph alignments. Genotypes at predefined SV sites were subsequently inferred using vg call, restricting genotype inference to alleles represented in the graph. The resulting multi-sample VCF was filtered for genotype completeness and minor allele frequency, and high-quality SVs were retained for subsequent association analyses. Structural variants were functionally annotated using SnpEff v5.4c (Cingolani, 2022). A custom SnpEff database was built from the B73 RefGen_v4 assembly and the corresponding annotation. SVs were annotated to determine their genomic context and predicted functional consequences, including intergenic, upstream/downstream, intronic, exonic and coding-sequence effects.

**Genome-wide association analyses based on small variants and structural variants**

Small variants- and structural variants (SV)-based GWAS were performed independently using Fast3VmrMLM (Wang et al., 2025). Small variants, comprising SNPs and short insertions/deletions (<30 bp), were analyzed as a single marker class, whereas structural variants >30 bp were analyzed separately. Small variants were filtered for MAF ≥ 0.05, marker missingness ≤ 0.05 and individual missingness ≤ 0.10. For SVs, insertions and deletions ≥30 bp were retained and filtered using MAF ≥ 0.05, marker missingness ≤ 0.20 and individual missingness ≤ 0.10. Population structure was accounted for using principal components derived from the genotype data. Association analyses were conducted with a 20-kb scanning window (svrad = 2 × 10⁴), svpal = 1 × 10⁻⁵ and a minimum LOD score of 3. GWAS was conducted independently for 22 agronomic traits and 91 flavor-related metabolites using the small-variant and SV datasets, resulting in four sets of association analyses: agronomic small-variant GWAS, agronomic SV-GWAS, metabolic small-variant GWAS and metabolic SV-GWAS. Significant QTNs identified by Fast3VmrMLM were retained for downstream analyses. Because the same genomic marker could be associated with multiple traits or detected in more than one GWAS dataset, association records were collapsed by variant class and marker identity to obtain a non-redundant set of QTL markers while retaining information on all associated traits and GWAS datasets.

**Parental genotype comparison at trait-associated loci**

To assess potential allelic complementation between the parental lines of Suyunuo 1, genotypes of Tongxi 5 and Hengbai 522 were compared at significant trait-associated loci. Significant QTL markers were matched to the genome-wide genotype dataset by marker identifier, chromosome and physical position, and parental genotypes were retrieved and converted to explicit diploid allele states according to the corresponding reference and alternative alleles. Loci were classified as identical, differentiated or missing between the two parents, and only loci with non-missing genotype calls in both lines were retained for comparison. The proportion of differentiated loci was calculated relative to the number of comparable QTLs. Parental differentiation at trait-associated loci was interpreted as potential genetic evidence for heterosis through allelic complementation.

**Colocalization of agronomic and metabolic QTLs**

To identify genomic regions potentially contributing to coordinated variation in agronomic performance and flavor-related metabolism, physical colocalization was assessed between agronomic and metabolic QTN signals. For each chromosome, all agronomic–metabolic QTN pairs separated by ≤250 kb were identified based on their physical coordinates. Pairs for which both QTNs reached the significance threshold were classified as significant agronomic–metabolic colocalizations, and significant QTN pairs separated by ≤100 kb were further designated as high-confidence colocalized pairs. To delineate broader candidate regions, QTNs participating in agronomic–metabolic colocalized pairs were clustered chromosome-wise. Adjacent participating QTNs separated by ≤250 kb were merged into the same candidate QTL region. For each region, the numbers of associated agronomic traits, metabolites and contributing QTN markers were summarized. Candidate regions containing both small-variant- and SV-based association signals were classified as jointly supported by the two variant classes. A hotspot score was calculated from the number of agronomic traits and metabolites represented within each region and was used to prioritize genomic regions showing extensive agronomic–metabolic connectivity. Because these analyses were based on physical overlap or proximity between independently detected GWAS signals, the resulting regions were interpreted as candidate colocalized QTLs reflecting shared genetic architecture.

Abrusán, G., Grundmann, N., DeMester, L. and Makalowski, W. (2009) TEclass—a tool for automated classification of unknown eukaryotic transposable elements. *Bioinformatics* **25**, 1329-1330.

Bailey, T.L., Williams, N., Misleh, C. and Li, W.W. (2006) MEME: discovering and analyzing DNA and protein sequence motifs. *Nucleic Acids Research* **34**, W369-W373.

Chen, J., Tang, Y., Yao, L., Wu, H., Tu, X., Zhuang, L. and Qi, Z. (2019) Cytological and molecular characterization of Thinopyrum bessarabicum chromosomes and structural rearrangements introgressed in wheat. *Molecular Breeding* **39**, 146.

Chen, S., Zhou, Y., Chen, Y. and Gu, J. (2018) fastp: an ultra-fast all-in-one FASTQ preprocessor. *Bioinformatics* **34**, i884-i890.

Chen, Y., Song, W., Xie, X., Wang, Z., Guan, P., Peng, H., Jiao, Y., Ni, Z., Sun, Q. and Guo, W. (2020) A Collinearity-Incorporating Homology Inference Strategy for Connecting Emerging Assemblies in the Triticeae Tribe as a Pilot Practice in the Plant Pangenomic Era. *Molecular Plant* **13**, 1694-1708.

Cheng, H., Concepcion, G.T., Feng, X., Zhang, H. and Li, H. (2021) Haplotype-resolved de novo assembly using phased assembly graphs with hifiasm. *Nature Methods* **18**, 170-175.

Cingolani, P. (2022) Variant Annotation and Functional Prediction: SnpEff. In: *Variant Calling: Methods and Protocols* (Ng, C. and Piscuoglio, S. eds), pp. 289-314. New York, NY: Springer US.

Danecek, P., Auton, A., Abecasis, G., Albers, C.A., Banks, E., DePristo, M.A., Handsaker, R.E., Lunter, G., Marth, G.T., Sherry, S.T., McVean, G., Durbin, R. and Genomes Project Analysis, G. (2011) The variant call format and VCFtools. *Bioinformatics* **27**, 2156-2158.

Delcher, A.L., Bratke, K.A., Powers, E.C. and Salzberg, S.L. (2007) Identifying bacterial genes and endosymbiont DNA with Glimmer. *Bioinformatics* **23**, 673-679.

Dobin, A., Davis, C.A., Schlesinger, F., Drenkow, J., Zaleski, C., Jha, S., Batut, P., Chaisson, M. and Gingeras, T.R. (2013) STAR: ultrafast universal RNA-seq aligner. *Bioinformatics* **29**, 15-21.

Du, P., Zhuang, L., Wang, Y., Yuan, L., Wang, Q., Wang, D., Dawadondup, Tan, L., Shen, J., Xu, H., Zhao, H., Chu, C. and Qi, Z. (2016) Development of oligonucleotides and multiplex probes for quick and accurate identification of wheat and Thinopyrum bessarabicum chromosomes. *Genome* **60**, 93-103.

Dudchenko, O., Batra, S.S., Omer, A.D., Nyquist, S.K., Hoeger, M., Durand, N.C., Shamim, M.S., Machol, I., Lander, E.S., Aiden, A.P. and Aiden, E.L. (2017) De novo assembly of the Aedes aegypti genome using Hi-C yields chromosome-length scaffolds. *Science* **356**, 92-95.

Durand, N.C., Robinson, J.T., Shamim, M.S., Machol, I., Mesirov, J.P., Lander, E.S. and Aiden, E.L. (2016a) Juicebox Provides a Visualization System for Hi-C Contact Maps with Unlimited Zoom. *Cell Systems* **3**, 99-101.

Durand, N.C., Shamim, M.S., Machol, I., Rao, S.S.P., Huntley, M.H., Lander, E.S. and Aiden, E.L. (2016b) Juicer Provides a One-Click System for Analyzing Loop-Resolution Hi-C Experiments. *Cell Systems* **3**, 95-98.

Ellinghaus, D., Kurtz, S. and Willhoeft, U. (2008) LTRharvest, an efficient and flexible software for de novo detection of LTR retrotransposons. *BMC Bioinformatics* **9**, 18.

Flynn, J.M., Hubley, R., Goubert, C., Rosen, J., Clark, A.G., Feschotte, C. and Smit, A.F. (2020) RepeatModeler2 for automated genomic discovery of transposable element families. *Proceedings of the National Academy of Sciences* **117**, 9451-9457.

Goel, M., Sun, H., Jiao, W.-B. and Schneeberger, K. (2019) SyRI: finding genomic rearrangements and local sequence differences from whole-genome assemblies. *Genome Biology* **20**, 277.

Gu, W., Yu, D., Guan, Y., Wang, H., Qin, T., Sun, P., Hu, Y., Wei, J. and Zheng, H. (2020) The dynamic transcriptome of waxy maize (Zea mays L. sinensis Kulesh) during seed development. *Genes & Genomics* **42**, 997-1010.

Guan, D., McCarthy, S.A., Wood, J., Howe, K., Wang, Y. and Durbin, R. (2020) Identifying and removing haplotypic duplication in primary genome assemblies. *Bioinformatics* **36**, 2896-2898.

Gurevich, A., Saveliev, V., Vyahhi, N. and Tesler, G. (2013) QUAST: quality assessment tool for genome assemblies. *Bioinformatics* **29**, 1072-1075.

Haas, B.J., Salzberg, S.L., Zhu, W., Pertea, M., Allen, J.E., Orvis, J., White, O., Buell, C.R. and Wortman, J.R. (2008) Automated eukaryotic gene structure annotation using EVidenceModeler and the Program to Assemble Spliced Alignments. *Genome Biology* **9**, R7.

Hickey, G., Heller, D., Monlong, J., Sibbesen, J.A., Sirén, J., Eizenga, J., Dawson, E.T., Garrison, E., Novak, A.M. and Paten, B. (2020) Genotyping structural variants in pangenome graphs using the vg toolkit. *Genome Biology* **21**, 35.

Hu, J., Wang, Z., Sun, Z., Hu, B., Ayoola, A.O., Liang, F., Li, J., Sandoval, J.R., Cooper, D.N., Ye, K., Ruan, J., Xiao, C.-L., Wang, D., Wu, D.-D. and Wang, S. (2024) NextDenovo: an efficient error correction and accurate assembly tool for noisy long reads. *Genome Biology* **25**, 107.

Huerta-Cepas, J., Forslund, K., Coelho, L.P., Szklarczyk, D., Jensen, L.J., von Mering, C. and Bork, P. (2017) Fast Genome-Wide Functional Annotation through Orthology Assignment by eggNOG-Mapper. *Molecular Biology and Evolution* **34**, 2115-2122.

Jain, C., Rhie, A., Zhang, H., Chu, C., Walenz, B.P., Koren, S. and Phillippy, A.M. (2020) Weighted minimizer sampling improves long read mapping. *Bioinformatics* **36**, i111-i118.

Kato, A. (1999) Air drying method using nitrous oxide for chromosome counting in maize. *Biotechnic & Histochemistry* **74**, 160-166.

Kim, D., Paggi, J.M., Park, C., Bennett, C. and Salzberg, S.L. (2019) Graph-based genome alignment and genotyping with HISAT2 and HISAT-genotype. *Nature Biotechnology* **37**, 907-915.

Lagesen, K., Hallin, P., Rødland, E.A., Stærfeldt, H.-H., Rognes, T. and Ussery, D.W. (2007) RNAmmer: consistent and rapid annotation of ribosomal RNA genes. *Nucleic Acids Research* **35**, 3100-3108.

Li, H. (2018) Minimap2: pairwise alignment for nucleotide sequences. *Bioinformatics* **34**, 3094-3100.

Li, H. (2023) Protein-to-genome alignment with miniprot. *Bioinformatics* **39**, btad014.

Li, H. and Durbin, R. (2009) Fast and accurate short read alignment with Burrows–Wheeler transform. *Bioinformatics* **25**, 1754-1760.

Li, H., Handsaker, B., Wysoker, A., Fennell, T., Ruan, J., Homer, N., Marth, G., Abecasis, G., Durbin, R. and Genome Project Data Processing, S. (2009) The Sequence Alignment/Map format and SAMtools. *Bioinformatics* **25**, 2078-2079.

Lowe, T.M. and Eddy, S.R. (1997) tRNAscan-SE: A Program for Improved Detection of Transfer RNA Genes in Genomic Sequence. *Nucleic Acids Research* **25**, 955-964.

Luo, J., He, C., Yan, S., Jiang, C., Chen, A., Li, K., Zhu, Y., Gui, S., Yang, N., Xiao, Y., Wu, S., Zhang, F., Liu, T., Wang, J., Huang, W., Yang, Y., Wang, H., Yang, W., Li, W., Zhuo, L., Fernie, A.R., Zhan, J., Wang, L. and Yan, J. (2024) A metabolic roadmap of waxy corn flavor. *Molecular Plant* **17**, 1883-1898.

Malinsky, M., Matschiner, M. and Svardal, H. (2021) Dsuite - Fast D-statistics and related admixture evidence from VCF files. *Molecular Ecology Resources* **21**, 584-595.

Manni, M., Berkeley, M.R., Seppey, M. and Zdobnov, E.M. (2021) BUSCO: Assessing Genomic Data Quality and Beyond. *Current Protocols* **1**, e323.

Marçais, G., Delcher, A.L., Phillippy, A.M., Coston, R., Salzberg, S.L. and Zimin, A. (2018) MUMmer4: A fast and versatile genome alignment system. *PLOS Computational Biology* **14**, e1005944.

Martin, S.H., Davey, J.W. and Jiggins, C.D. (2015) Evaluating the Use of ABBA–BABA Statistics to Locate Introgressed Loci. *Molecular Biology and Evolution* **32**, 244-257.

McKenna, A., Hanna, M., Banks, E., Sivachenko, A., Cibulskis, K., Kernytsky, A., Garimella, K., Altshuler, D., Gabriel, S., Daly, M. and DePristo, M.A. The Genome Analysis Toolkit: A MapReduce framework for analyzing next-generation DNA sequencing data. **20**, 1297-1303.

Nawrocki, E.P. and Eddy, S.R. (2013) Infernal 1.1: 100-fold faster RNA homology searches. *Bioinformatics* **29**, 2933-2935.

Ou, S., Chen, J. and Jiang, N. (2018) Assessing genome assembly quality using the LTR Assembly Index (LAI). *Nucleic Acids Research* **46**, e126-e126.

Ou, S. and Jiang, N. (2018) LTR_retriever: A Highly Accurate and Sensitive Program for Identification of Long Terminal Repeat Retrotransposons  *Plant Physiology* **176**, 1410-1422.

Ou, S., Su, W., Liao, Y., Chougule, K., Agda, J.R.A., Hellinga, A.J., Lugo, C.S.B., Elliott, T.A., Ware, D., Peterson, T., Jiang, N., Hirsch, C.N. and Hufford, M.B. (2019) Benchmarking transposable element annotation methods for creation of a streamlined, comprehensive pipeline. *Genome Biology* **20**, 275.

Pertea, M., Pertea, G.M., Antonescu, C.M., Chang, T.-C., Mendell, J.T. and Salzberg, S.L. (2015) StringTie enables improved reconstruction of a transcriptome from RNA-seq reads. *Nature Biotechnology* **33**, 290-295.

Quinlan, A.R. and Hall, I.M. (2010) BEDTools: a flexible suite of utilities for comparing genomic features. *Bioinformatics* **26**, 841-842.

Ramani, V., Deng, X., Qiu, R., Lee, C., Disteche, C.M., Noble, W.S., Shendure, J. and Duan, Z. (2020) Sci-Hi-C: A single-cell Hi-C method for mapping 3D genome organization in large number of single cells. *Methods* **170**, 61-68.

Rauluseviciute, I., Riudavets-Puig, R., Blanc-Mathieu, R., Castro-Mondragon, Jaime A., Ferenc, K., Kumar, V., Lemma, R.B., Lucas, J., Chèneby, J., Baranasic, D., Khan, A., Fornes, O., Gundersen, S., Johansen, M., Hovig, E., Lenhard, B., Sandelin, A., Wasserman, Wyeth W., Parcy, F. and Mathelier, A. (2024) JASPAR 2024: 20th anniversary of the open-access database of transcription factor binding profiles. *Nucleic Acids Research* **52**, D174-D182.

Rhie, A., Walenz, B.P., Koren, S. and Phillippy, A.M. (2020) Merqury: reference-free quality, completeness, and phasing assessment for genome assemblies. *Genome Biology* **21**, 245.

Sirén, J., Monlong, J., Chang, X., Novak, A.M., Eizenga, J.M., Markello, C., Sibbesen, J.A., Hickey, G., Chang, P.-C., Carroll, A., Gupta, N., Gabriel, S., Blackwell, T.W., Ratan, A., Taylor, K.D., Rich, S.S., Rotter, J.I., Haussler, D., Garrison, E. and Paten, B. Pangenomics enables genotyping of known structural variants in 5202 diverse genomes. *Science* **374**, abg8871.

Stanke, M., Diekhans, M., Baertsch, R. and Haussler, D. (2008) Using native and syntenically mapped cDNA alignments to improve de novo gene finding. *Bioinformatics* **24**, 637-644.

Tempel, S. (2012) Using and Understanding RepeatMasker. In: *Mobile Genetic Elements: Protocols and Genomic Applications* (Bigot, Y. ed) pp. 29-51. Totowa, NJ: Humana Press.

Wang, J., Chen, Y., Shu, G., Zhao, M., Zheng, A., Chang, X., Li, G., Wang, Y. and Zhang, Y.-M. (2025) Fast3VmrMLM: A fast algorithm that integrates genome-wide scanning with machine learning to accelerate gene mining and breeding by design for polygenic traits in large-scale GWAS datasets. *Plant Communications* **6**.

Wolff, J., Rabbani, L., Gilsbach, R., Richard, G., Manke, T., Backofen, R. and Grüning, B.A. (2020) Galaxy HiCExplorer 3: a web server for reproducible Hi-C, capture Hi-C and single-cell Hi-C data analysis, quality control and visualization. *Nucleic Acids Research* **48**, W177-W184.

Xu, S., Hu, E., Cai, Y., Xie, Z., Luo, X., Zhan, L., Tang, W., Wang, Q., Liu, B., Wang, R., Xie, W., Wu, T., Xie, L. and Yu, G. (2024) Using clusterProfiler to characterize multiomics data. *Nature Protocols* **19**, 3292-3320.

Xu, Z. and Wang, H. (2007) LTR_FINDER: an efficient tool for the prediction of full-length LTR retrotransposons. *Nucleic Acids Research* **35**, W265-W268.

Yan, S., Huang, W., Gao, J., Fu, H. and Liu, J. (2018) Comparative metabolomic analysis of seed metabolites associated with seed storability in rice (Oryza sativa L.) during natural aging. *Plant Physiology and Biochemistry* **127**, 590-598.

Yang, J., Lee, S.H., Goddard, M.E. and Visscher, P.M. (2011) GCTA: A Tool for Genome-wide Complex Trait Analysis. *The American Journal of Human Genetics* **88**, 76-82.

Zhao, X., Chai, Q., Yin, W., Fan, H., He, W. and Zhao, C. (2025) Organic fertilizer substitution altered the waxy maize grain quality and aroma volatiles formation by the integrated transcriptomic and metabolomic analyses. *Frontiers in Plant Science* **Volume 16 - 2025**.

Zhou, Z.-W., Yu, Z.-G., Huang, X.-M., Liu, J.-S., Guo, Y.-X., Chen, L.-L. and Song, J.-M. (2022) GenomeSyn: a bioinformatics tool for visualizing genome synteny and structural variations. *Journal of Genetics and Genomics* **49**, 1174-1176.

Zhu, M., Du, P., Zhuang, L., Chu, C., Zhao, H. and Qi, Z. (2017) A simple and efficient non-denaturing FISH method for maize chromosome differentiation using single-strand oligonucleotide probes. *Genome* **60**, 657-664.
